# Structural characterization of pyruvic oxime dioxygenase, a key enzyme of heterotrophic nitrification

**DOI:** 10.1101/2024.10.08.617318

**Authors:** Shuhei Tsujino, Yusuke Yamada, Miki Senda, Akihiko Nakamura, Toshiya Senda, Taketomo Fujiwara

## Abstract

Nitrification by heterotrophic microorganisms is an important part of the nitrogen cycle in the environment. The enzyme responsible for the core function of heterotrophic nitrification is pyruvic oxime dioxygenase (POD). POD is a non-heme Fe(II)-dependent enzyme that catalyzes the dioxygenation of pyruvic oxime to produce pyruvate and nitrite. To analyze the catalytic mechanism of POD, the crystal structure of POD from *Alcaligenes faecalis* (AfPOD) was determined at 1.76 Å resolution. The enzyme is a homo-tetramer and the subunit structure is homologous to those of class II aldolases, in particular a zinc-dependent L-fuculose-1-phosphate aldolase. The active site of the subunit is located at the bottom of a cleft formed with an adjacent subunit. The iron ion at the active site is coordinated by three histidines and three water molecules in an octahedral geometry. The putative oxygen tunnel was connected between the active site and the central cavity of the tetramer. The N-terminal region of AfPOD, which is essential for catalytic activity, is disordered in the crystal. Structure prediction with AlphaFold2 combined with mutational experiments suggested that the disordered N-terminal region adopts an α-helix conformation and participates in the formation of the active site. The catalytic mechanism of the dioxygenase reaction by POD is discussed on the basis of the molecular docking model.

**IMPORTANCE:** Our knowledge of nitrification has increased considerably in recent decades with the discovery of new nitrifying microorganisms and the characterization of their biochemical processes. Some heterotrophic bacteria and fungi are known to show nitrification activities, but the molecular mechanisms had been poorly understood. Here, we performed a structural characterization of POD, a key enzyme in heterotrophic nitrification that produces nitrite from ammonia using pyruvic oxime as an intermediate. Structural and enzymatic analyses revealed that POD is a unique dioxygenase with features such as an aldolase backbone, an N-terminal α-helix, and an oxygen tunnel. Our results provide insights not only into the molecular mechanisms but also into the design of specific inhibitors of heterotrophic nitrification.

## INTRODUCTION

Nitrification, the catabolic oxidation of ammonia, is a microbiological process that is essential to the global nitrogen cycle. In addition, understanding the diversity and metabolic mechanisms of nitrifying microorganisms would lead to applications such as preventing nitrogen loss from fertilized cropland and ammonia removal technology for wastewater treatment (**1, 2**).

It has long been known that heterotrophic microorganisms, including various species of bacteria and fungi, perform dissimilatory ammonia oxidation to nitrite or nitrate (**3**), termed heterotrophic nitrification. Heterotrophic nitrification is thought to contribute significantly to the nitrogen cycle in the environment due to its species richness and abundance (**4**). Bacteria of the genera *Alcaligenes* and *Pseudomonas* are known to perform heterotrophic nitrification, which produces nitrite from ammonia using pyruvate oxime (PO) as an intermediate metabolite (**5–7**). Ono *et al*. (**8**) reported the presence of pyruvate oxime dioxygenase (POD), a dioxygenase that converts PO to nitrite, from *Alcaligenes faecalis.* However, the molecular and catalytic properties of POD have not been fully investigated since its discovery.

Therefore, we purified POD from *A. faecalis* (AfPOD) and showed that AfPOD is a metalloenzyme that requires Fe(II) ion for catalytic activity. AfPOD is a homotetramer, and its amino acid sequence is classified as a class II aldolase (PF00596) (**9**). We also confirmed that POD is involved in heterotrophic nitrifying activity (**9, 10**). Sequence homology searches using genome databases revealed *pod* genes in many types of bacteria in the phyla Proteobacteria, Actinobacteria, and fungi in the phylum Ascomycota (**9, 11**). While heterotrophic nitrifying activities have not been reported from many microbial species with *pod* genes, this result suggests that heterotrophic nitrifying microbes with the *pod* gene contribute more to the nitrogen cycle in the environment than previously thought.

Here we report the first X-ray crystal structure of a novel dioxygenase, AfPOD, at 1.76 Å resolution. While the overall structure of AfPOD is similar to the class II aldolase with the zinc ion in the active site, AfPOD has the iron ion in the active site. Although the N-terminal 18 amino acids are critical for the enzyme activity of AfPOD, the N-terminal region was disordered in the crystal. Structure prediction using AlphaFold2 suggests that the disordered N-terminal region of AfPOD adopts an α-helix conformation and is involved in the formation of the active site. The catalytic mechanism of the dioxygenase reaction by POD was discussed based on the predicted substrate-binding structure obtained by molecular docking simulation.

## MATERIALS AND METHODS

### Preparation of purified samples of AfPOD, AfPODdN18 and BWPOD

AfPOD was overexpressed in the *Escherichia coli* cells and purified by two chromatographic steps: anion-exchange chromatography on DEAE-Toyopearl 650 M gel (Tosoh, Tokyo, Japan) and gel filtration chromatography on Sephacryl S-200 (Cytiva, Marlborough, MA), as described previously (**9**). Sample purity was confirmed by SDS-PAGE (**12**). The construction of the overexpression system of the variant enzyme (AfPODdN18) lacking the N-terminal 18 residues and the purification of the recombinant were performed by a similar procedure as for AfPOD with some modifications as described in the Supplementary section. The construction of the expression plasmid, overexpression and purification of the recombinant protein of POD from *Bradyrhizobium* sp. WSM3983 (BWPOD), which was successfully crystallized in the early stage of the project, is also described in the Supplementary section.

### Crystallization of AfPOD, AfPODdN18, and BWPOD

Crystallization conditions were initially screened using PEGIon/PEGIon2 (Hampton Research, Aliso Viejo, CA), Crystal Screen Cryo and 2 Cryo (Hampton Research), Wizard Screens I and II (Rigaku, Tokyo, Japan), and PEGsII (Qiagen, Hilden, Germany) with a Protein Crystallization System 2 (PXS2) at the Structural Biology Research Center, High Energy Accelerator Research Organization, Japan (**13**). The obtained crystallization conditions were further optimized manually using the sitting-drop vapor diffusion method. However, since AfPOD is gradually deactivated even at 0°C due to iron dissociation from the active site (**9**), the AfPOD in the crystal was an inactive form. Therefore, reconstitution with Fe(II) and subsequent crystallization of AfPOD were performed in an anaerobic chamber with mixed gas (N_2_/H_2_, 96:4, v/v) and Pd catalyst to maintain anaerobic conditions (**14**). Reconstitution of AfPOD with Fe(II) was performed in the reservoir solution supplemented with 1 mM FeSO_4_. AfPOD crystals were obtained at 20°C in 36% (v/v) PEG400, 0.2 M calcium acetate, 0.1 M HEPES (pH 7.5) with 5.0 mg ml^−1^ of purified AfPOD, by the sitting-drop vapor-diffusion method (**Supplementary Fig. S1a**). In addition, crystallization of the variant enzyme (AfPODdN18) lacking the N-terminal 18 residues was performed under aerobic conditions with Mn(II) ions. AfPODdN18 crystals were obtained at 20°C in 20 % (w/v) PEG 3350, 1 mM manganese chloride, 0.2 M sodium tartrate (pH 7.3) with 5.0 mg ml^−1^ of purified AfPODdN18, by the sitting-drop vapor-diffusion method (**Supplementary Fig. S1b**). Both crystals were successfully obtained under each condition as shown in **Supplementary Fig. S1**. The BWPOD crystals were obtained at 20°C in 1.56 M ammonium sulfate and 25% (v/v) glycerol with 10 mg ml^−1^ of purified BWPOD by the sitting-drop vapor-diffusion method as described in the supplementary section.

### Data collection and structure determination

Crystals of AfPOD and BWPOD were cryoprotected with the corresponding crystallization solution. Crystals of AfPODdN18 were cryoprotected with the crystallization solution containing 30% (v/v) glycerol. All crystals were frozen in liquid nitrogen. The X-ray diffraction data sets of BWPOD were collected at –178°C using a beam wavelength of 2.7 Å at BL-1A of Photon Factory (Tsukuba, Japan) and processed using XDS (**15**). Phase determination and initial model construction of BWPOD were performed using CRANK2 (**16**) by the native SAD method (**17**). The X-ray diffraction data sets of AfPOD and AfPODdN18 were collected at –178°C using a beam wavelength of 0.98 Å at Photon Factory’s BL-17A. The diffraction data sets were processed using AutoPROC (**18**) due to the anisotropy of the crystals. Molecular replacement was performed with MOLREP using the structure of BWPOD as a search model (**19**). Model building and crystallographic refinement were performed using COOT, REFMAC5, and BUSTER (**20–22**).

The metal species that bind to the active site were identified using anomalous difference Fourier maps calculated from diffraction data sets with X-rays on the shorter and longer wavelength sides of their absorption edges. All crystallographic figures were generated using PyMOL (Schrödinger Inc, New York, NY).

### Site-directed mutagenesis of AfPOD

Site-directed mutagenesis of the AfPOD was performed by the technical application of PCR. A set of oligonucleotide primers, in which the corresponding original amino residue was replaced by the mutation, was amplified by using the AfPOD expression vector with C-terminal His_6_-tag as a template. After treatment with the restriction enzyme *Dpn*I to digest the template DNA, the PCR product was introduced into the *E. coli* BL21-CodonPlus(DE3). The PCR product was cyclized by homologous recombination between the 5’– and 3’-regions in the host cells, generating the strain for overexpression of the variant AfPOD. After confirmation of the nucleotide sequence, the resulting strain was cultured aerobically in the 2×YT medium (30 mL). Expression was performed in the same manner as for the wild-type AfPOD. The variant AfPOD was purified by His-tag purification using His-Spin Protein Miniprep (Zymo Research, Irvine, CA) according to the manufacturer’s protocol. Primers used for the PCR amplification are listed in **Supplementary Table S1**.

### Homology analysis and structure prediction

Alignment analysis and conserved amino acid estimation were performed using Clustal Omega (**23**), MEGA X (**24**), and ConSurf server (**25**). Comparison with protein structures deposited in the PDB was performed using the DALI server (**26**). Structure prediction of the N-terminal part of AfPOD was performed using AlphaFold2 (**27**) as implemented by the ColabFold server (**28**).

### Enzymatic analysis

POD activity was determined by measuring the rate of nitrite production in the assay solution containing 20 mM Tris-HCl buffer (pH 8.0), 1.0 mM sodium ascorbate, 100 μM FeSO_4_, and 1.0 mM pyruvic oxime. The concentration of nitrite was measured spectrophotometrically using a diazo-coupling method (**29**). Each reaction was run in triplicate. GraphPad Prism 9 (GraphPad Prism Software Inc, San Diego, CA) was used for statistical data analysis. Pyruvic oxime was synthesized according to Quastel *et al.* (**30**). Protein concentration was measured using a BCA protein assay kit (Pierce, Rockford, IL) with bovine serum albumin as the standard. Spectroscopic analysis in the visible region was performed in a 1 cm light path cuvette using a UV-2600 spectrophotometer (Shimadzu, Kyoto, Japan). All chemicals used in the experiments were of the highest quality commercially available.

### Docking simulation

In the present study, co-crystallization or soaking experiments did not yield the structure of AfPOD bound to pyruvic oxime. We attempted to dock the substrate with the AfPOD structure that was generated by removing the binding water molecules on Fe ion from the crystal structure (8IL8) and adding hydrogens at pH 7 using PyMOL. The docking simulation was performed using Gnina (https://github.com/gnina/gnina), a convolutional neural network (CNN) molecular docking program, with an accurate line option (**31**). The EcFucA (4FUA) was aligned with AfPOD using the three catalytic histidine residues. Search regions were defined using the autobox_ligand option with the superimposed phosphoglycohydroxamic acid (PGH).

## RESULTS

### Conserved amino acids analysis of AfPOD

The internal region of 178 residues of AfPOD, excluding the N-terminal 32 residues and the C-terminal 51 residues, has been annotated as a class II aldolase (Pfam00596) and showed a 20.7 % amino acid sequence identity with L-fuculose-1-phosphate aldolase (FucA), whose crystal structure has already been solved (**9**). Interestingly, a homology search using the KEGG genome database revealed that many class II aldolase genes with unknown functions shared extremely high sequence homology (E-value < e-60) with AfPOD. Molecular phylogenetic analysis has shown that these proteins form a monophyletic group that is distinct from other class II aldolases such as FucA, and recombinant experiments have confirmed their POD activity (**11**). The amino acid sequence alignment of AfPOD, BWPOD and *E. coli* FucA (EcFucA) is shown in **Fig. 1**. The result of a conservation analysis of amino acid residues, which was conducted using the sequences of all 452 PODs identified in the KEGG genome database, is also shown in the figure. Since the His-triads (His119, His121, His183, AfPOD numbering) corresponding to the zinc-binding motif of EcFucA, are completely conserved, they are predicted to coordinate the Fe(II) ion in the active site of the POD (**9, 32**). Highly conserved regions, such as Glu45-Ser55, Gly72-Ala76, Asn98-His104, and Trp155-Gly165 (AfPOD numbering), were found in the PODs. In addition, highly conserved amino acids such as Lys8 and Met19 were found in the N-terminal region, which is not observed in class II aldolases and is a unique feature of PODs.

**Fig. 1.**
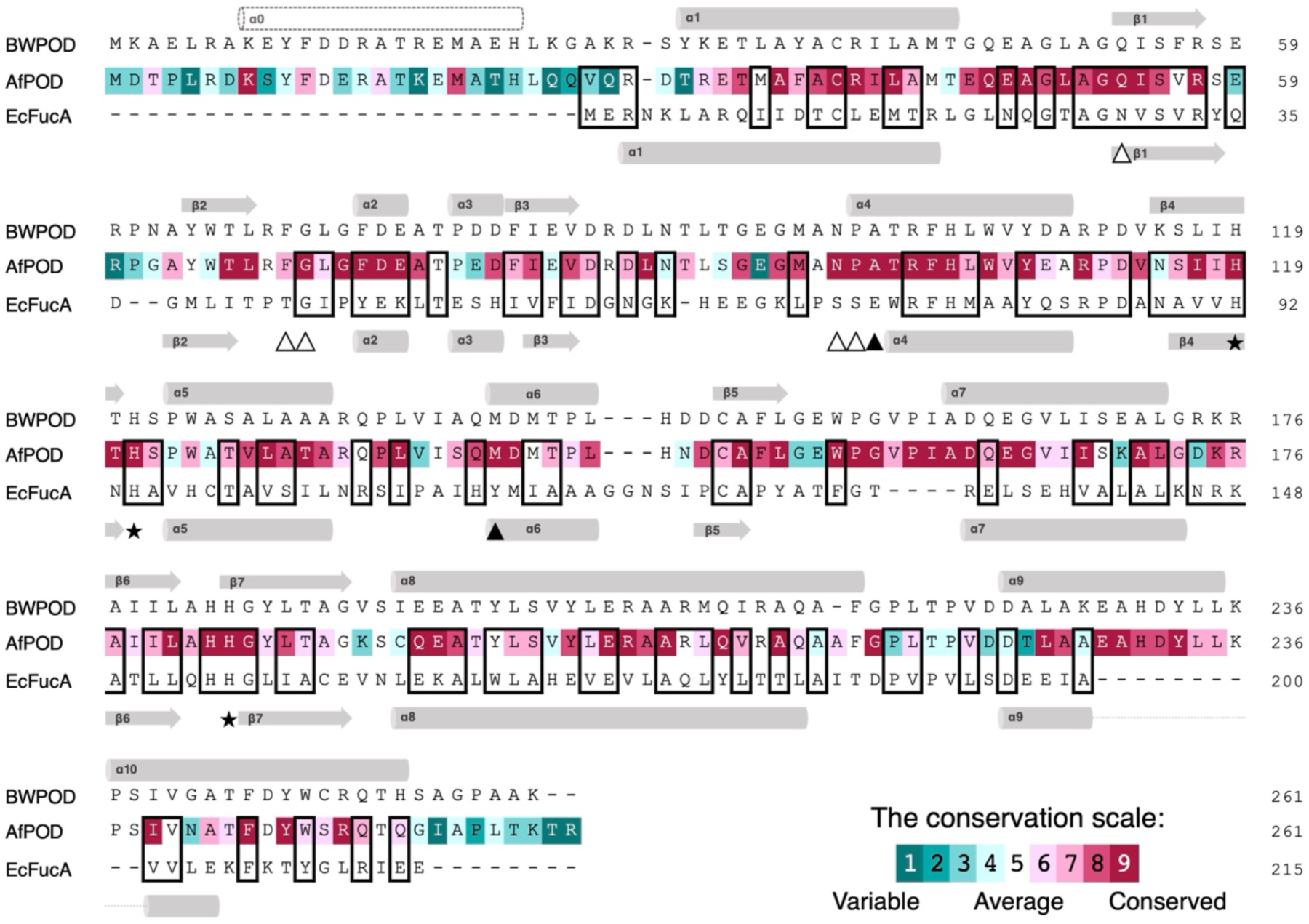
Sequence alignment of AfPOD, BWPOD and EcFucA. The amino acid sequences of AfPOD, BWPOD and EcFucA were aligned, and conserved amino acids are shown in boxes. Arrows and bars indicate α-helix and β-sheet structures, respectively. The amino acid sequence of AfPOD is indicated according to the conservation scale in the PODs. The amino acids of EcFucA for metal binding (His triad), substrate recognition, and catalytic residues are indicated by stars (★), white triangle (△), and black triangle (▴), respectively.

### Overall structure of AfPOD

The crystal structures of AfPOD and AfPODdN18 were determined at resolutions of 1.76 Å and 1.55 Å, respectively (**Table 1**). Although crystal structure analysis of the AfPOD-substrate complex was attempted by the soaking method under anaerobic conditions, no electron density was observed for the substrate molecule.

**Table 1.**
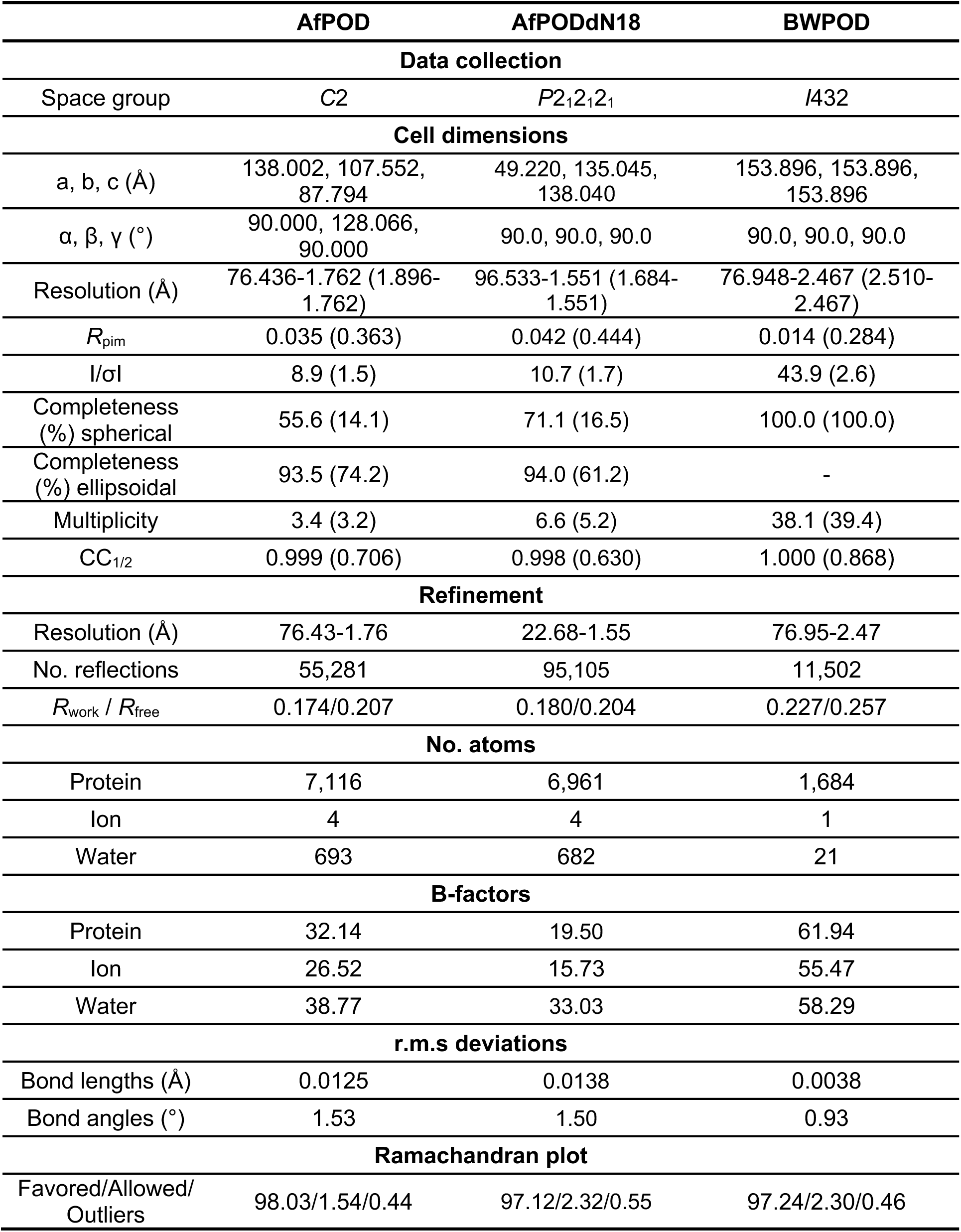
Data collection and refinement statistics.

The final structure model of the AfPOD tetramer contains 920 amino acid residues, 693 water molecules, and four iron ions. The N-terminal region (Met1-Arg28) and the C-terminal region (Lys259-Arg261) of each subunit showed no apparent electron density. The AfPOD monomer consists of a seven-stranded anti-parallel β-sheet (β5↑, β6↑, β7↓, β4↑, β1↓, β2↑, β3↓) flanked by two α-helices on one side (α4 and α7) and eight α-helices (α1, α2, α3, α5, α6, α8, α9, α10) on the opposite side (**Fig. 2a**). AfPOD has a tetrameric structure with approximate dimensions of 60 x 40 x 80 Å^3^ (**Fig. 2b**). The central cavity of the tetramer has the 18 Å diameter opening on the C-terminal side and is closed by the last turn of α8 on the opposite side (**Fig. 2c**). The exposed surface area of the tetramer is approximately 45,000 Å^2^, of which 1,522 Å^2^ and 7 Å^2^ are buried at the interface between the A and B subunits and that between the A and C subunits, respectively. The solvation free energy (ΔG) gain calculated by PISA (**33**) is –19.2 kcal/mol between the A and B, the potential number of hydrogen bonds is 16, and the number of salt bridges is 10; while between the A and C, these values are –0.2 kcal/mol, 0, and 0, respectively.

**Fig. 2.**
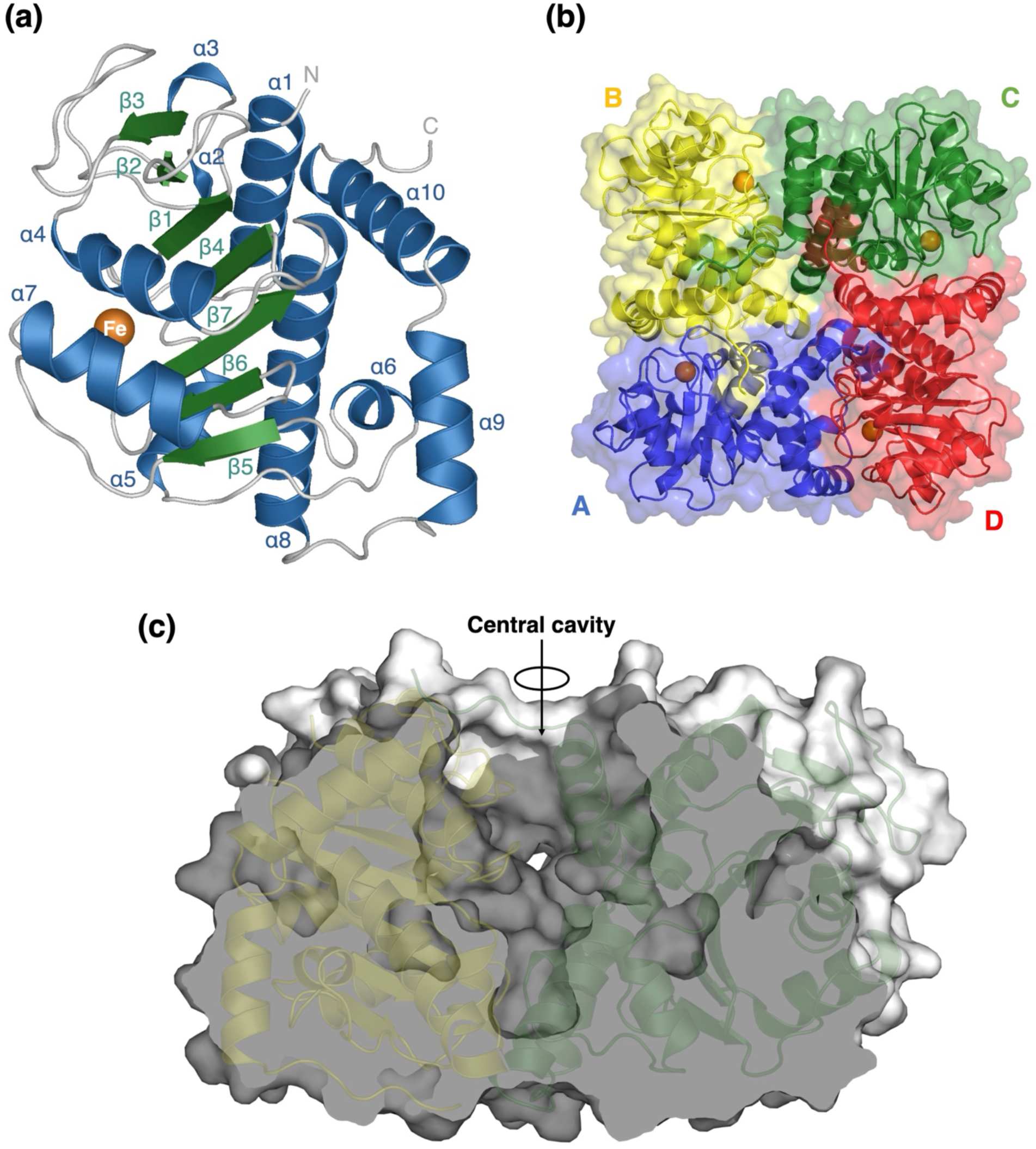
Monomeric and tetrameric structures of AfPOD. (**a**) Ten α-helices (blue) and seven β-sheets (green) in the monomeric structure of AfPOD. (**b**) Top view of the entire AfPOD molecule along the fourfold axis. Each subunit is indicated by a single color (A, blue; B, yellow; C, green; D, red). (**c**) Cross-sectional view of the entire AfPOD molecule. The iron ions are shown in orange.

### Putative active site structure

The anomalous difference Fourier maps revealed that the metal ion coordinated by three His residues was confirmed to be an iron ion, as expected from a previous biochemical analysis (**Supplementary Fig. S2a,b**). In subunit A, the iron ion is located at a depth of about 8 Å at the bottom of an active site pocket with an opening of about 16 Å long and 8 Å wide at the boundary to the adjacent subunit B (**Fig. 3a**). The opening is constricted by the side chains of Phe69, Tyr233’, and Ile239’. Here, an attached label (’) to an amino acid indicates that it is derived from a neighboring subunit molecule. Imidazole nitrogen atoms of His119, His121, and His183 and oxygen atoms of three water molecules (W1, W2 and W3) coordinate the iron ion, forming an octahedral arrangement. The geometric parameters of the ligands are summarized in **Supplementary Table S2**. The side chains of Gln53, Phe103, His104, and Glu164 were located around the iron center. These amino acid residues (His119, His121, His183, Gln53, Phe103, His104, and Glu164) are completely conserved in different PODs (**Fig. 1**). The α1-β1 loop (Glu45-Ala51), which is highly conserved among PODs, is also located near the iron center and is fixed by hydrogen bonding between the main-chain carbonyl oxygen of Ala51 and the iron-coordinated water 2 at distances of 2.5 Å. Interestingly, the side chain of Glu47 in the mobile loop shows a different conformation in subunits A, C and B, D (**Supplementary Fig. S3**), which may affect the transport of oxygen molecules to the active site (see below).

**Fig. 3.**
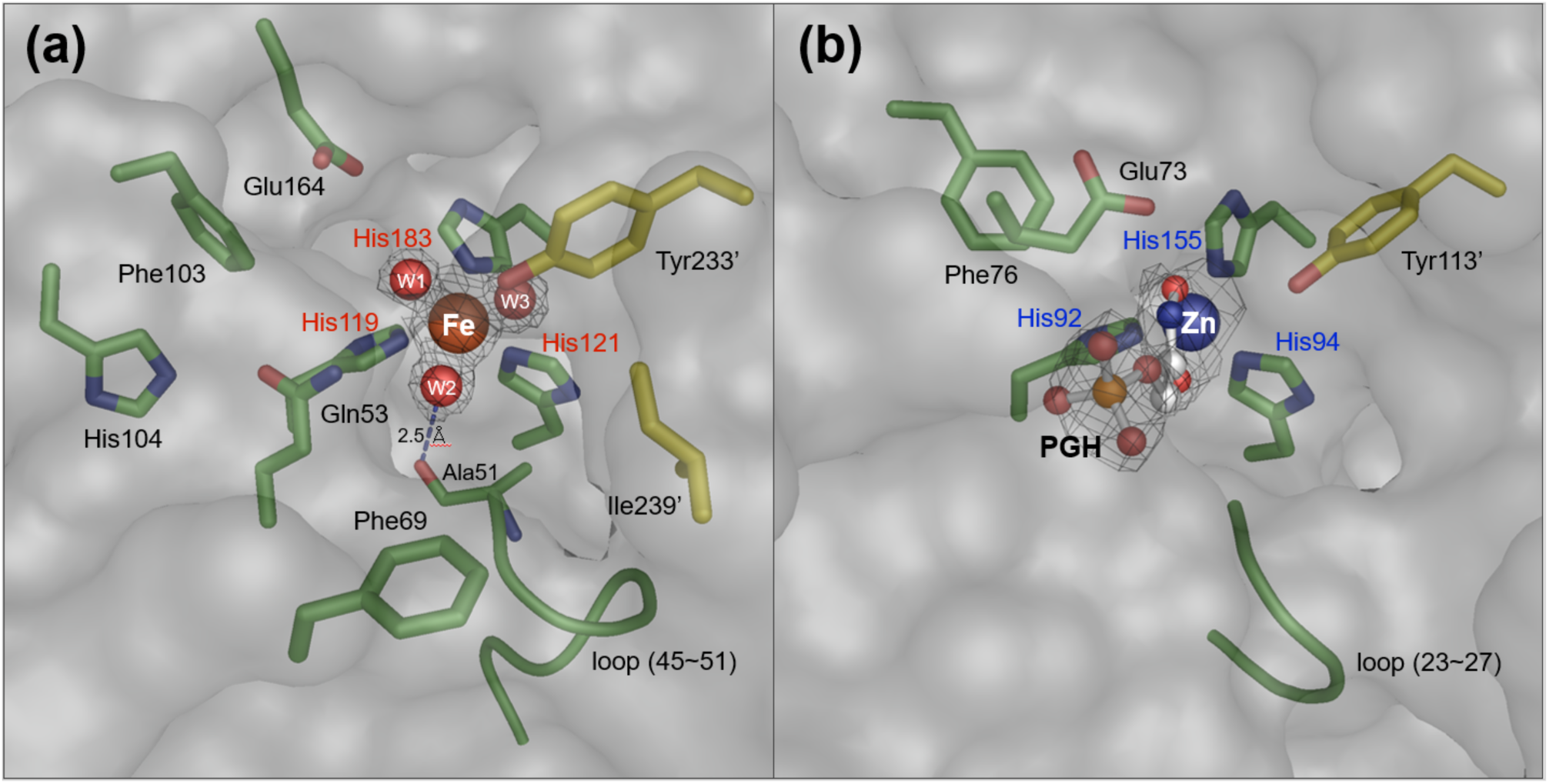
Comparison of the active site structures of AfPOD and EcFucA. The active site structure of AfPOD (8IL8) (**a**) was compared with that of EcFucA (4FUA) (**b**). The side chains and loop structure around the His triad are shown. Electron density maps around the iron and ligands are also shown. Residues shown in green are from subunit A, and the residues shown in yellow with an attached label (’) are from the neighboring subunit B.

### Mutagenesis analysis of AfPOD

Of the amino acids highly conserved between PODs, six amino acid residues located near the putative active site, Gln53, Phe103, His104, Glu164, Phe69, and Tyr233’, were substituted by alanine and the activity of each variant was assayed (**Table 2**). The catalytic constant (*k*_cat_) of the POD activity of the Q53A, H104A, E164A, and F69A variants was reduced to about 3-5% of the value of the wild-type enzyme, and was further reduced to 0.2-0.3% in the F103A and Y233A variants. The Michaelis constant (K_m_) values for the Q53A, F103A and H104A variants were approximately 0.1 to 0.2 mM in comparison to the wild-type enzyme (0.622 mM), indicating an enhancement of substrate affinity due to the mutations. All three variants, Q53A, F103A, and H104A, showed similar catalytic changes in increased substrate affinity, indicating that the Gln53, Phe103, and His104 residues are not only structurally but also functionally coordinated. In contrast, substitutions of Phe69 or Tyr233’ to alanine significantly reduced substrate affinity. Tyr233’ is conserved in all the PODs as an aromatic amino acid (Tyr, Trp, or Phe) (**11**). The side chains of Phe69 and Tyr233′ are 8.3 and 12.1 Å away from the iron center, respectively, and are not expected to interact directly with the substrate molecule bound to the iron center. The result suggests that the hydrophobic region formed by the side chains of Tyr233’ and Phe69 in the vicinity of the active site is important for the enzymatic activity of POD.

**Table 2.** Kinetic parameters of wild-type and variant AfPODs. The values in parentheses of *k_cat_*/K*_m_* and *k_cat_* indicate the relative value to WT. N.D. = Not Detected.

| | $k_{cat}/K_m$ (s<sup>-1</sup> M<sup>-1</sup>) | $k_{cat}$ (s<sup>-1</sup>) | $K_m$ (mM) |
| --- | --- | --- | --- |
| WT | $7.7 \times 10^3$ (100) | $4.797 \pm 0.169$ (100) | $0.622 \pm 0.083$ |
| dN18 | – | N.D. | N.D. |
| Q53A | $1.7 \times 10^3$ (22.0) | $0.162 \pm 0.008$ (3.4) | $0.096 \pm 0.018$ |
| F103A | $9.0 \times 10$ (1.2) | $0.008 \pm 0.001$ (0.2) | $0.089 \pm 0.022$ |
| H104A | $1.2 \times 10^3$ (16.0) | $0.210 \pm 0.013$ (4.4) | $0.170 \pm 0.037$ |
| E164A | $2.9 \times 10^2$ (3.7) | $0.270 \pm 0.020$ (5.6) | $0.943 \pm 0.240$ |
| F69A | 5.9 (0.1) | $0.246 \pm 0.018$ (5.1) | $41.509 \pm 9.984$ |
| Y233A | 0.3 (0.003) | $0.012 \pm 0.001$ (0.3) | $42.247 \pm 16.705$ |
| K8E | 9.2 (0.1) | $0.632 \pm 0.066$ (13.2) | $68.766 \pm 20.299$ |
| E229K | $2.1 \times 10^3$ (27.6) | $4.580 \pm 0.098$ (95.5) | $2.150 \pm 0.172$ |
| K8E/E229K | $6.9 \times 10^2$ (9.0) | $1.762 \pm 0.109$ (36.7) | $2.538 \pm 0.568$ |

### Structure prediction of the disordered N-terminal region

Since an AfPODdN18 variant lacking the N-terminal 18 amino acid residues showed no POD activity (**Table 2**), the N-terminal region should play an important role in the POD activity. However, this region was disordered in the AfPOD crystal; therefore, no structural information on the N-terminal region was obtained. The structure of AfPODdN18 crystallized in the coexistence of manganese was determined as a tetramer containing 918 amino acid residues, 682 water molecules, and four metal ions. The N-terminal regions (Met19-Arg28) and C-terminal regions (Lys259-Arg261) of each subunit showed no apparent electron densities. The overall structure of AfPODdN18 is nearly identical to the AfPOD with a Cα root mean square deviation (RMSD) of 0.3 Å (845 Cα atoms). The anomalous difference Fourier maps suggested that the electron density near the His triad in the AfPODdN18 is not a manganese ion (**Supplementary Fig. S2c,d**). Nickel or cobalt ions may have been bound to the His triad, rather than manganese, during overexpression or purification.

AlphaFold2, a high accurate structure prediction software, has been reported to be useful for predicting the structure of disorder regions (**34**). Therefore, we used AlphaFold2 implemented on the ColabFold server to predict the structure of the entire length of the AfPOD, including the disordered region. The predicted structure showed a high agreement with the AfPOD crystal structure with a Cα RMSD of 0.5 Å (720 Cα atoms, **Supplementary Fig. S4**). In the predicted structure, the N-terminus of the AfPOD forms an α-helix (Lys8-His22), and the C-terminus is further extended towards an adjacent subunit (**Fig. 4**). The N-terminal α-helix, which is designated as helix α0, of an adjacent subunit covers the putative the active site pocket, and hydrophobic interactions between the side chains of Thr16’, Met19’ and Leu23’ on the α-helix and those of Phe69, Tyr233’ and Ile239’ at the pocket opening were expected. It is reasonable to assume that helix α0 is a part of the active site.

**Fig. 4.**
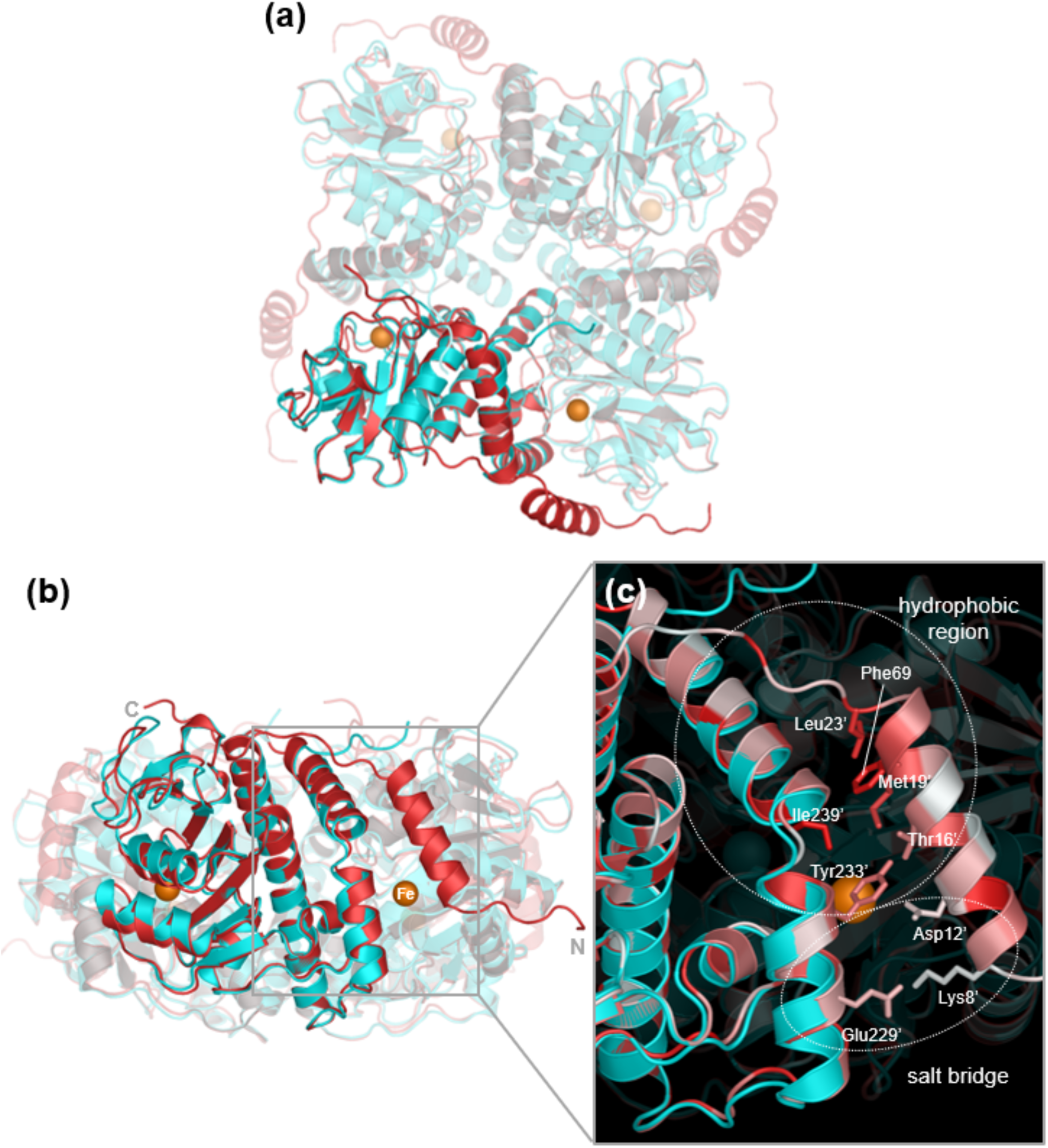
Comparison of the AfPOD crystal structure and the structure predicted by AlphaFold2. (**a**) Top and (**b**) side views of the structures. (**c**) The crystal structures (cyan) and their predicted structures (red) have been superimposed. The zoomed views show the active site of subunit A covered by an N-terminal α0-helix extending from the adjacent subunit B. The amino acid residues of subunit B are marked with an attached label. The hydrophobicity and hydrophilicity of the amino acid side chains are color-coded in red and white, respectively.

The formation of a salt bridge by electrostatic interactions between the side chains of Lys8’ and Glu229’, which are highly conserved in POD, was also suggested (**Figs. 1 and 4**). Three variants, K8E, E229K and K8E/E229K, were generated and their POD activities were measured to assess the importance of the salt bridge (**Table 2**). The POD activity of the K8E variant was significantly reduced in both substrate affinity and catalytic activity, and then the specificity constant (*k*_cat_/K_m_) was reduced to 0.1% of that of the wild-type (WT) enzyme. In the E229K variant, the catalytic constant was not affected, but the substrate affinity was reduced to about 30% of that of the WT enzyme. Interestingly, while the introduction of the K8E mutation into the wild-type enzyme resulted in a significant decrease in substrate affinity, the K8E mutation into the E229K variant had only a small effect on substrate affinity. This result can be interpreted as the substrate affinity and catalytic constant, which were significantly lowered in the K8E variant, were restored by the further introduction of the E229K mutation. Since Lys8 is the only basic amino acid near the active site, replacing it with an acidic amino acid could disrupt the structure of the active site by preventing salt bridge formation. It was suggested that the substitution of Glu229 in K8E variant for basic Lys restores the structure of the active site by reforming the salt bridge between the two residues, resulting in some recovery of enzyme activity.

### Putative oxygen pathway

A putative oxygen pathway extending from the central cavity of the AfPOD tetramer to the active site has been identified. A 14 Å long tunnel with a 3.5 Å diameter opening at the central cavity extends along the interface between the subunit molecules to the iron center (**Fig. 5**). Notably, the α1-β1 loop (Glu45-Ala51) forms part of the tunnel in subunits B and D. However, the tunnel is blocked by a conformational change of the Glu47 in subunits A and C, which probably prevents the passage of oxygen molecules (**Fig. 5a,b**). Since the deeper part of the tunnel and the active site pocket are hydrophobic, oxygen molecules may be recruited to the pocket by diffusion due to their hydrophobic nature (**35**). The structural difference of the α1-β1 loop suggested that the oxygen supply to the active site pocket of POD was controlled by the movement of the α1-β1 loop. The entrance to the active site pocket opened in the region proximal to the side chains of Phe69, Tyr233’, and Ile239’. In the closed state, where the N-terminal α-helix covers the pocket opening, this space is further covered by the hydrophobic side chains of Thr16’, Met19’, and Leu23’, making this space highly hydrophobic and suitable as an oxygen binding site.

**Fig. 5.**
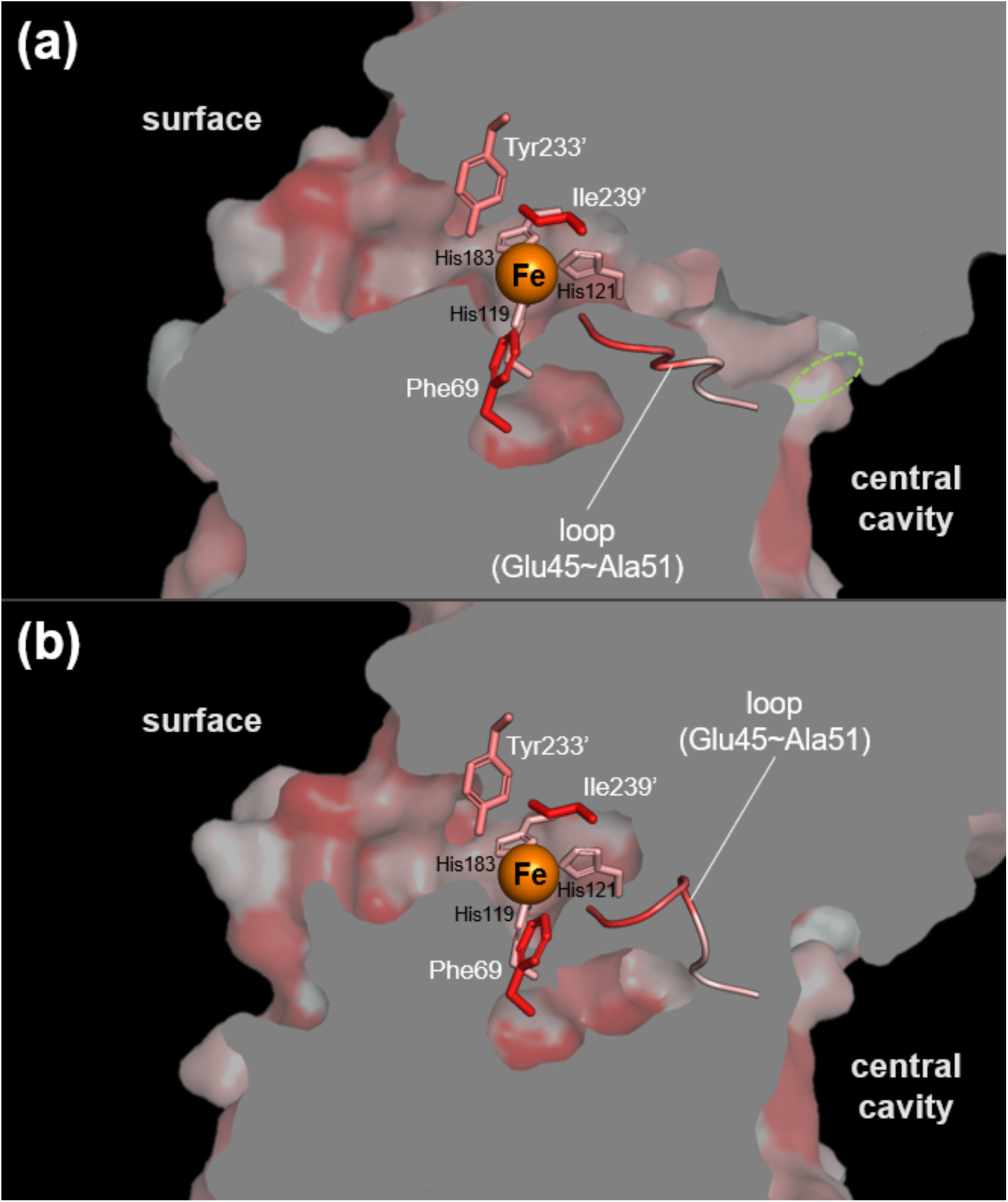
Cross-section view of the putative oxygen tunnel. The hydrophobicity of the tunnel interior leading to the active site of the subunit A (**a**) and that of the subunit D (**b**) are color-coded (red, hydrophobic; white, hydrophilic). The entrance of a tunnel leading to the active site is circled in green. His triad (His119, 121, 183) and side chains around the active site pocket (Phe69, Tyr233’ and Ile239’) and loop structure (Glu45∼Ala51) are shown in the figure.

### Docking simulation of the active site with pyruvic oxime

Since the crystal structure of AfPOD in the bound state with pyruvate oxime was not obtained in this experiment, we attempted to estimate the substrate binding structure of AfPOD by docking simulation. Docking with the Gnina program yielded a total of 20 structures with a CNN pose score greater than 0.4. (**Supplementary Table S3**). In the three structures with the highest pose scores (model 1∼3), the carbonyl oxygen and the oxime oxygen of the pyruvate oxime were each located at the positions of two of the three coordinated water molecules of iron. In the most likely model 1 structure, the carbonyl and oxime oxygens were located at positions W1 and W2, respectively, and the carbonyl carbon, C2-carbon, oxime nitrogen, and oxime oxygen of the pyruvate oxime formed a six-membered ring structure with Fe(II) (**Supplementary Fig. S5**). The oxime oxygen of the pyruvic oxime was positioned at a distance of approximately 3 Å, allowing hydrogen bonding, with the carbonyl oxygen of the Ala51 main chain and the amido nitrogen of the Glu53 side chain. The carboxyl oxygen of the substrate molecule was also predicted to form hydrogen bonds with the amido nitrogen of the Glu53 side chain and the carbonyl oxygen of the Glu164 side chain. The conserved amino acid residues Phe103, His104, Phe69, and Tyr233, which have been shown to be essential or strongly involved in POD activity, did not interact directly with the substrate molecule in the docking model.

## DISCUSSION

While several cytochrome P450-type enzymes are known to catalyze the monooxygenation of oxime (aldoxime) compounds to form nitriles (**36**), POD is the only dioxygenase that uses oxime compounds as substrates to produce nitrite. As expected from the results of the sequence homology analysis, AfPOD showed remarkable structural homology to class II aldolases, which are zinc-dependent lyases. A structural similarity search using the DALI server revealed that AfPOD is structurally similar to class II aldolases (COG0235) with known functions, such as FucA, 5-deoxyribose disposal aldolase (DrdA), L-ribulose-5-phosphate 4-epimerase (AraD), and L-rhamnulose-1-phosphate aldolase (Rua). FucA showed the highest structural homology to AfPOD among them. These enzymes require a zinc ion for their activity. In addition, AfPOD showed a structural similarity to Mn(II)-dependent 3-hydroxy-2-methylpyridine-4,5-dicarboxylate decarboxylase (HMPDdc) (**37-40**) (**Supplementary Table S4**). No dioxygenases showed structural similarity to AfPOD. Superposition of the AfPOD subunit with EcFucA (PDB code: 1FUA), HMPDdc (2Z7B), DrdA (6BTG), and EcAraD (1JDI) (**Fig. 6**) showed that the seven-stranded anti-parallel β-sheet structure flanked by α-helices from both sides of the β-sheet is well conserved among them. PODs have additional sequences at the N-terminus consisting of 20-30 amino acid residues that are not present in class II aldolases, corresponding to the disordered region of AfPOD in its crystal structure.

**Fig. 6.**
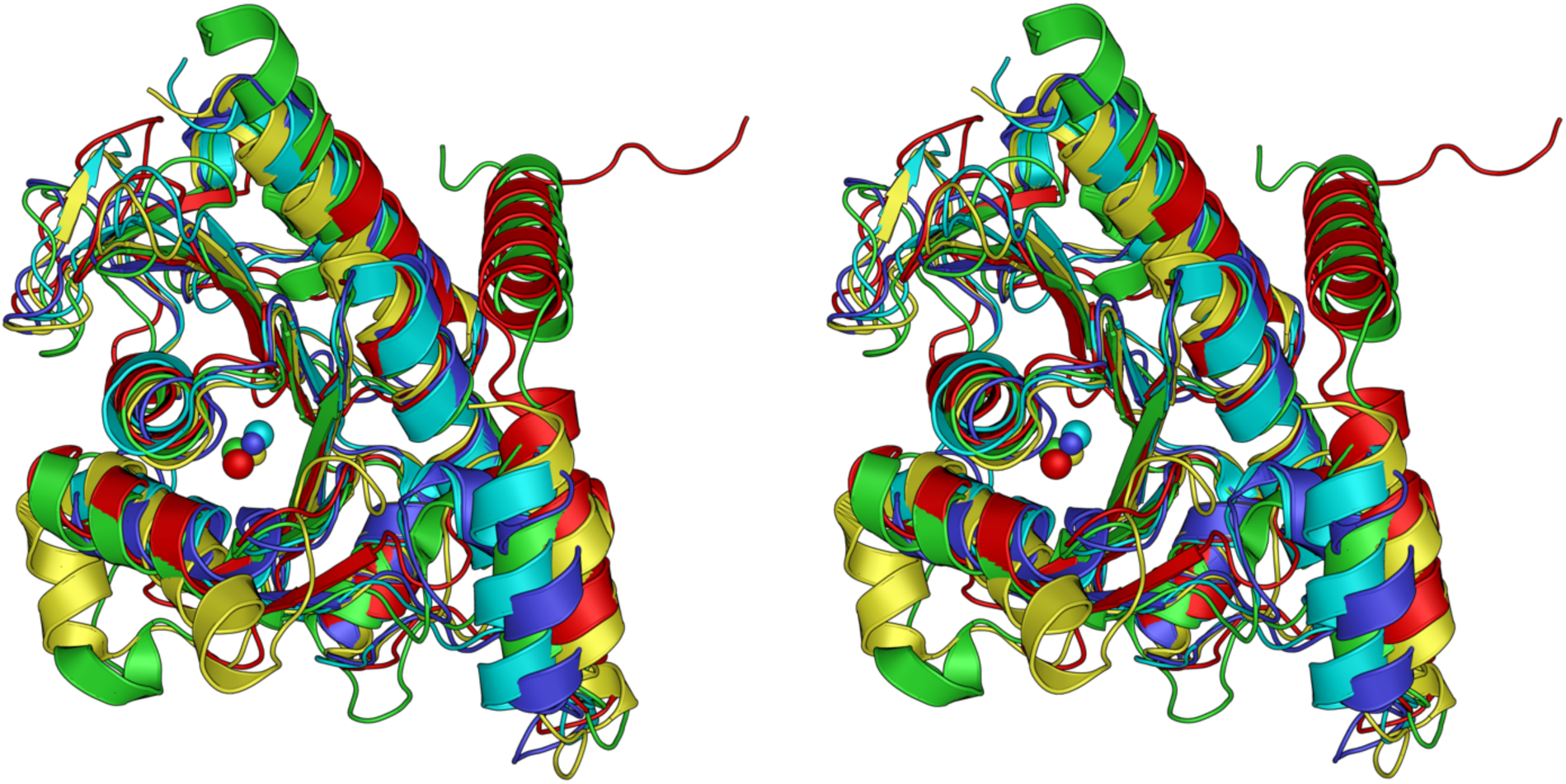
Stereoview diagram of the superposition of AfPOD with the homologous enzymes. The monomeric structures of the protein groups with particularly high similarity (Z-score ≥ 20.0) are superimposed based on the structural homology search on the DALI server. AfPOD (PDB code: 8IL8), EcFucA (1FUA), HMPDdc (2Z7B), DrdA (6BTG), and EcAraD (1JDI) are shown in red, blue, green, cyan, and yellow, respectively.

The active site of AfPOD has an iron ion that has an octahedral coordination sphere with six ligands: three His residues and three water molecules (**Supplementary Table S2**). On the other hand, with a few exceptions such as cysteine dioxygenase, the non-heme Fe(II)-dependent dioxygenases have an iron center with two His residues and one Glu (or Asp) residue (**41**, **42**). One possible explanation is that POD is the enzyme that was evolved by metal substitution using the protein folding common to the class II aldolases. The scenario that an oxidoreductase (dioxygenase) evolved from a lyase (aldolase) by replacing the zinc ion with the iron ion is unique in terms of the molecular evolution of metalloenzymes and should be further investigated.

The reaction mechanism of FucA has already been analyzed, and important catalytic residues have been identified. In **Fig. 3**, the structure of the active site of the AfPOD was compared with that of the EcFucA bound to the inhibitor phosphoglycolohydroxamic acid (PGH) (PDB code: 4FUA). The aldol cleavage of L-fuculose-1-phosphate by FucA is initiated by the coordination of the carbonyl oxygen at the C2 position and the hydroxyl oxygen at the C3 position of L-fuculose in the open chain state to the zinc ion to stabilize the cis-enediolate structure. Cleavage of the C3-C4 bond is then promoted by deprotonation of the C4 hydroxyl group by the Tyr113’ and that of the C3 hydrogen by the Glu73 (**43**). Two amino acids of EcFucA, Glu73 and Tyr113’, which are essential for the catalytic function of FucA, are not conserved in the AfPOD sequence (**Fig. 1**) (**44**). The amino acid residues, Asn29, Thr43, Gly44, Ser71, and Ser72, which are involved in substrate recognition of EcFucA by hydrogen bonding to the phosphate group of PGH, were also not conserved in the active site pocket of AfPOD. However, Glu164 and Tyr233’ in the active site of the AfPOD are structurally conserved with EcFucA; Glu164 and Tyr233’ are located at the positions corresponding to Glu73 and Tyr113’ of EcFucA, respectively (**Fig. 3**). Phe76 in EcFucA, which was proposed to interact with protonated Glu73 side chains, was conserved as Phe103 in AfPOD (**Figs. 1** and **3**) (**32**). The α1-β1 loop (Asn23-Ala27) of EcFucA, which corresponds to the α1-β1 loop (Glu47-Ala51) of AfPOD, is also highly mobile and has been suggested to be essential for substrate binding by the point mutagenesis experiment conducted on EcFucA (**45**). The main chain carbonyl oxygen of Ala27 has been proposed to be involved in the rearrangement of the hydrogen bond network upon substrate binding, leading to an induced fit at the active site of EcFucA (**43**). In the case of AfPOD, the movement of the α1-β1 loop may be involved in the regulation of O_2_ supply to the active site by switching of the oxygen tunnel open/closed (**Fig. 5**). In addition, the main chain carbonyl oxygen of the corresponding Ala51 of AfPOD is thought to promote the POD reaction by hydrogen bonding with the substrate molecule, as discussed below.

Based on the docking model 1 with the highest CNN pose score and the known catalytic mechanism of extradiol-type dioxygenase, the molecular mechanism of catalysis by POD was predicted as shown in **Fig. 7**. In the model 1 structure, the carbonyl and oxime oxygens of the pyruvic oxime are located at positions W1 and W2, respectively, forming a six-membered ring containing Fe(II) (**Supplementary Fig. S5**). In addition, the oxime oxygen of the substrate molecule is hydrogen-bonded to the carbonyl oxygen of the Ala51 main chain and the amide nitrogen of the Glu53 side chain, and the carboxyl oxygen is hydrogen-bonded to the amide nitrogen of the Glu53 side chain and the carbonyl oxygen of the Glu164 side chain, respectively. Aldol reactions catalyzed by class 2 aldolases proceed via the tautomeric enol form of the substrate molecule, which is stabilized by the interaction of the carbonyl oxygen with an electron-withdrawing zinc ion (**43**). In the pyruvic oxime molecule, the oxime group (=N-OH) is in tautomeric equilibrium with the nitroso group (-N=O), and the equilibrium is usually strongly biased toward the oxime-type structure. Within the six-membered ring in docking model 1, the equilibrium is predicted to shift toward the nitroso form due to the electron withdrawing effect of Fe(II) as a Lewis acid. In addition, the carbonyl oxygen of the main chain of Ala51 is close enough to form a hydrogen bond with the oxime oxygen and may contribute to the stabilization of the nitroso-form structure by accepting the oxime hydrogen. The shift in the oxime/nitroso tautomeric equilibrium and the deprotonation would lead the C2 carbon of the pyruvic oxime to an electron-rich state. O_2_ molecules entering the active site through the oxygen channel are expected to bind to the W3 position near the channel opening (**Fig. 3a**). A new carbon-oxygen bond is formed between the O_2_ and the C2 carbon, forming a six-membered transition state ring consisting of the C2 carbon, the nitroso nitrogen, the nitroso oxygen, two oxygen atoms, and an iron atom. Subsequent isomerization of the six-membered ring transition state, i.e., cleavage of the oxygen-oxygen bond and attack of the nitrogen by the oxygen atom on the iron, would result in the formation of nitrite and pyruvate.

**Figure 7.**
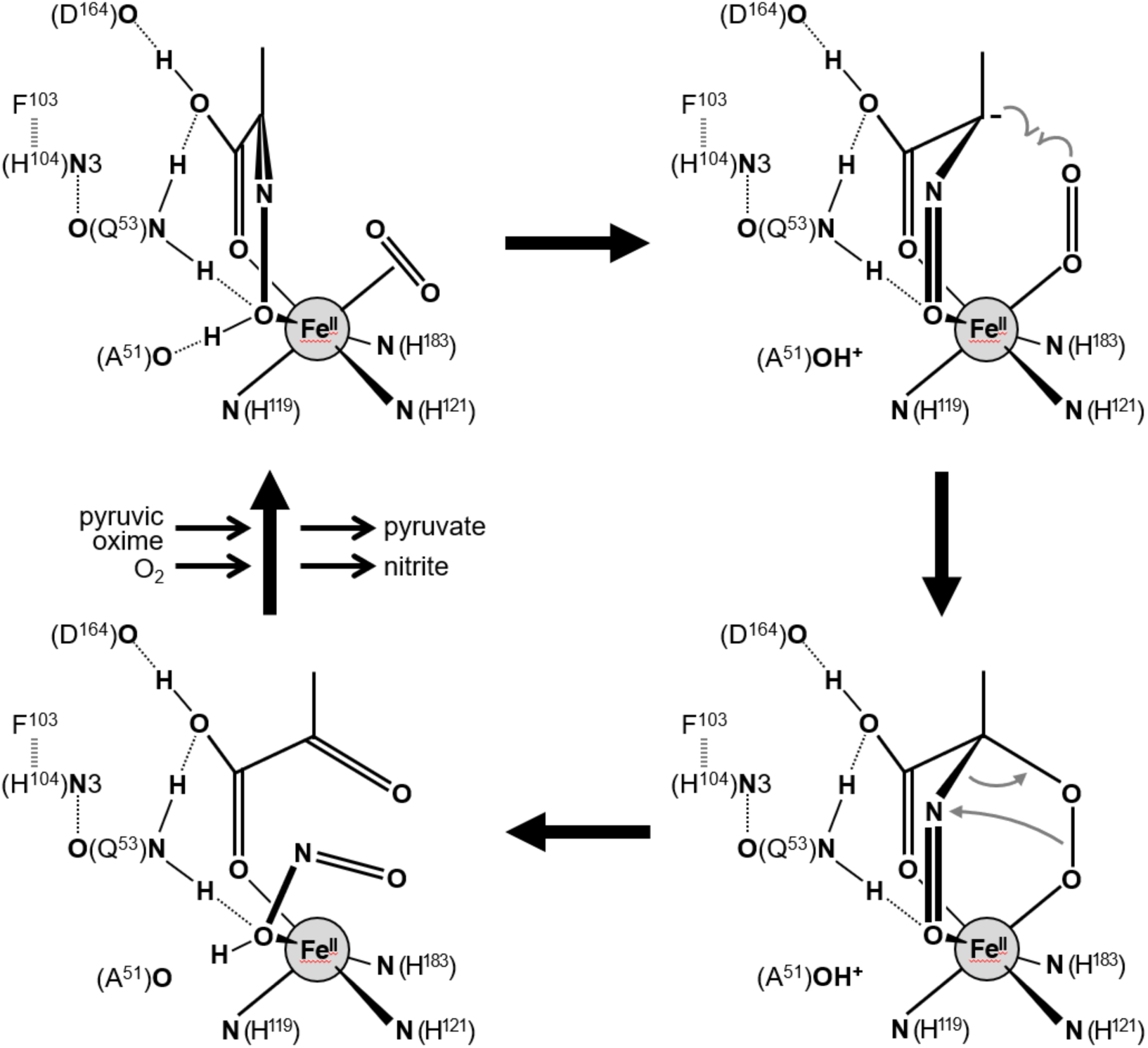
Proposed reaction mechanism of POD. Dotted line and thick dotted line show hydrogen bonding and π-π stacking interactions, respectively. See text for further details.

In the structure of the docking model 1, hydrogen bonds with the side chains of Gln53 and Asp164, and with the main chain of Ala51 are expected to contribute to the binding of the pyruvic oxime molecule to the active site of AfPOD. A significant decrease of the catalytic constant *k*_cat_ was observed in the Q53A and E164A variants compared to the wild-type enzyme. On the other hand, the substrate affinity increased in the Q53A variant, whereas it was little affected in the E164A variant (**Table 1**).

Interestingly, the F103A and H104A variants also exhibited the decreased catalytic activity and increased substrate affinity observed in the Q53A variant. The side chains of the three amino acid residues Gln53, Phe103, and His104 are located close to each other within the putative active site, and π-π stacking interactions between the aromatic rings of Phe103 and His104, and hydrogen bonds between the imidazole N3 nitrogen of His104 and the carbonyl oxygen of the Gln53 side chain were indicated by crystal structure analysis (**Fig. 3A**). In the presumed reaction mechanism shown in **Fig. 7**, the C2 carbon of the pyruvate oxime and the O_2_ molecule bonded to the W3 position would need to face each other in an appropriate configuration. These results suggest that the Phe103 and His104 side chains may indirectly promote the catalytic reaction by anchoring the Gln53 side chain in the appropriate position in the active site. In the Q53A mutant, the carbamoylethyl side chain is replaced by a small methyl side chain, which would facilitate the entry of the substrate molecule into the active site. However, it will be less likely to hold the substrate molecule in the appropriate spatial arrangement within the active site. The enzymatic properties observed in the F103A and H104A variants, which are similar to those of the Q53A variant, can be attributed to the inability to anchor Glu53 in its proper position.

The dN18 variant of AfPOD completely lost its enzymatic activity, indicating that the POD-specific N-terminal extension of about 30 amino acid residues, which is not present in other class 2 aldolases, is essential for catalytic activity (**Table 1**). Initially, we expected the flexibility of the region to cause disorder in the N-terminal sequence of AfPOD and BWPOD crystals, but the AlphaFold2 prediction gave an α-helix conformation in the sequence (**Fig. 4**). The active site pocket of subunit A was predicted to be covered by this N-terminal helix, designated α0, extending from subunit B. However, in the crystal it could not assume such a configuration due to steric hindrance with other molecules in the lattice and thus appeared to be observed as in the disordered state. Electrostatic interaction to form a salt bridge between the side chains of Lys8’ on the α0 helix and the Asp229’ on the α9 helix were also postulated to be involved in the “closed” state in which the α0 helix covers the active site pocket (**Table 1 and Fig. 4**). No entrance of the substrate to the active site was found when the α0 helix was in the closed state. Therefore, the α0 helix would have to switch to the “open” state for the substrate molecule to enter the active site. At present, it is most acceptable to assume that this conformational switching between the closed and the open state of the α0 helix plays an important role in the uptake of the substrate. When the α0 helix returns to the closed state after substrate uptake, a hydrophobic environment is expected to be formed in the active site pocket as shown in **Fig. 4**. This hydrophobic nature may contribute not only to the recruitment of O_2_ molecules by diffusion through the tunnel, but also to the formation of a suitable environment for the dioxygenase reaction in the active site pocket.

The marked decrease in substrate affinity observed in the F69A, Y233A, and K8E mutants may be due to some impairment in the formation of the hydrophobic environment in the active site (**Table 1**).

In this study, the crystal structure of POD from the heterotrophic nitrifying bacterium *A. faecalis* was determined. The structure of AfPOD and its homologous enzyme BWPOD were very similar to those of class II aldolases, whereas the zinc ion, which is the active center in other class II aldolases, was replaced by Fe(II) ion in the active state of AfPOD. Class II aldolases catalyze aldol reactions by shifting the keto-enol tautomeric equilibrium of the substrate molecule to the enol side through the electron-withdrawing effect of the zinc ion. In POD, the electron withdrawing action of Fe(II) ions is thought to shift the oxime-nitroso tautomeric equilibrium of the substrate molecule to the nitroso side, thereby promoting the reaction with Fe(II)-bound O_2_ and the removal of the nitroso group. An extended sequence at the N-terminus of POD, not present in other class II aldolases, was disordered in the crystal structure of AfPOD, but was inferred to form an α-helix and positioned to cover the active pocket by the AlphaFold2 structure prediction. The conformational switching of this α0 helix between the open and closed states is thought to be important for the catalytic process of POD. The catalytic mechanism discussed above should be verified based on the crystal structure of substrate-bound AfPOD. For example, the AfPOD variants with increased substrate affinity might be useful for preparing crystals of the substrate-binding enzyme. Furthermore, the conformational switching of the α0 helix of AfPOD could be directly confirmed by cryo-EM observations.

## CONCLUSIONS

AfPOD is a unique dioxygenase with high amino acid sequence similarity to class II aldolases such as EcFucA. We have determined the crystal structure of POD at 1.76 Å resolution, revealing a class II aldolase-like overall structure. The active site of AfPOD contains the Fe(II) that is coordinated by three His residues and three water molecules. While functionally important N-terminal residues were not observed in the crystal structure, a combination of the AlphaFold2 structure prediction and point mutation experiments suggested the critical function of the N-terminal (putative) α-helix in the catalytic reaction. In addition, the functional roles of amino acid residues around the iron center, and the migration pathway of oxygen molecules were inferred. The catalytic mechanism of the POD was discussed on the basis of the molecular docking with the pyruvic oxime. This finding raises the possibility that class II aldolases of unknown function, present in the genomes of many microorganisms, may function as dioxygenases and be responsible for diverse metabolic functions other than aldolases. The findings also provide a platform for the future design of nitrification inhibitors specific for heterotrophic nitrification.

### Date accessibility

The coordinates and the structure factor amplitudes for the structure of AfPOD, AfPODdN18, and BWPOD have been deposited in the Protein Data Bank (PDB) under accession codes 8IL8, 8IQA, and 8IX6.

### Author’s contributions

S.T.: conceptualization, data curation, formal analysis, investigation, methodology, validation, visualization, writing—original draft and writing—review and editing; Y.Y.: data curation, formal analysis, investigation, methodology, validation, writing—original draft and writing—review and editing; M.S.: methodology, investigation, and writing—review and editing; A.N.: investigation, methodology, writing— original draft and writing—review and editing; T.S.: methodology, supervision, and writing—review and editing; T.F.: conceptualization, funding acquisition, project administration, supervision, validation, writing—original draft and writing—review and editing.

All authors gave final approval for publication and agreed to be held accountable for the work performed therein.

## Acknowledgements

We thank the staff of the Structural Biology Research Center, Photon Factory, Institute of Materials Structure Science, High Energy Accelerator Research Organization for crystallization. We also thank Mr. Yuichi Azuma at the Shizuoka University for assistance in constructing the BWPOD expression vector. This work was supported by KAKENHI (17K00517, 21K12220, and 24K15258) from the Ministry of Education, Culture, Sports, Science and Technology, Japan, and Basis for Supporting Innovative Drug Discovery and Life Science Research (BINDS) from the Japan Agency for Medical Research and Development (AMED). An Amano Institute of Technology Scholarship also supported S.T. to continue the investigation.

## Supplementary materials

### Supplementary Methods

#### Construction of the AfPODdN18 expression system

A set of oligonucleotide primers, AfpoddN18f and AfpodR, was designed for the construction of the mutant AfPOD lacking the 18 N-terminal amino acid residues (AfPODdN18) expression vector.

Amplification was performed using KOD-plus DNA polymerase (Toyobo, Osaka, Japan) and the AfPOD expression vector (pAfPOD) as a template (**1**). The resulting PCR product was cloned into a pCR-blunt TOPO II vector (Invitrogen, Carlsbad, CA), yielding pCRAfPODdN18. After confirmation of the nucleotide sequence, the insert of pCRAfPODdN18 was digested with both *Nde*I and *Xho*I, and then cloned into the same restriction site of a pET21a+ vector (Novagen, Darmstadt, Germany), yielding the expression plasmid pAfPODdN18. The pAfPODdN18 plasmid was introduced into *E. coli* BL21-CodonPlus(DE3) (Agilent Technologies, Santa Clara, CA) to generatie the strain AfPdN18 for overexpression of AfPODdN18. Standard protocols used for DNA handling in *E. coli* were followed Sambrook and Russell (**2**). Primers used for the amplification are listed in **Supplementary Table S1**.

### Purification of AfPODdN18

The purification of AfPODdN18 was performed as described above with slight modification using AKTA FPLC (Cytiva). The strain AfPdN18 was cultured aerobically in the 2×YT medium (1 L) supplemented with 50 μg mL-1 ampicillin at 37°C with reciprocal shaking at 150 rpm. At a mid-exponential growth stage (OD600 = 0.6–0.8), isopropyl β-D-1-thiogalactopyranoside (IPTG) was added to the medium to reach 0.3 mM for induction of AfPODdN18. After incubation at 20°C with shaking at 150 rpm for 3 h, the cells were collected by centrifugation. Cultured AfPdN18 cells were suspended in 30 mL of 20 mM Tris-HCl (pH 8.0) containing 10 μM phenylmethylsulfonyl fluoride (PMSF) (buffer A) and disrupted by sonication. After removal of unbroken cells by centrifugation at 12,000 × g for 10 min, the supernatant obtained was centrifuged at 140,000 × g for 65 min. The soluble fraction thus obtained was applied to a HiTrap DEAE FF anion-exchange chromatography column (Cytiva) equilibrated with buffer A. The recombinant protein adsorbed on the column was eluted by a linear gradient generated from 100 mL each of buffer A and buffer A containing 0.4 M NaCl. The fractions confirmed by SDS-PAGE were collected and then concentrated by 30–50% saturated ammonium sulfate fractionation. The precipitate obtained was suspended in 500 μL of buffer A containing 0.25 M NaCl and then applied to a gel filtration chromatography column of Superdex 200 Increase 10/300 GL (Cytiva) equilibrated with the same buffer. The fractions confirmed by SDS-PAGE were collected and concentrated by ammonium sulfate fractionation as above. The resulting precipitate was suspended in 10 mM HEPES buffer (pH 7.0) and dialyzed three times against the same buffer. The purified protein was electrophoretically homogenized by SDS-PAGE and then used for crystallization. All purification experiments were carried out at low temperatures around 4°C.

### Construction of the BWPOD expression system and purification of the recombinant BWEPOD

The gene encoding BWPOD, whose nucleotide sequence was optimized according to an *E. coli* codon usage, was synthesized (GENEWIZ Inc, South Plainfield, NJ). The 777 bp fragment was cloned into an *Nde*I and *Xho*I site of a pET21a(+) vector, yielding the expression plasmid pBWPOD. The pBWPOD plasmid was introduced into *E. coli* BL21-CodonPlus(DE3) to generate strain BW01 for overexpression of BWPOD. The His_6_-tagged BWPOD was overexpressed in strain BW01 and purified using a Ni^2+^-chelating Sepharose column (Cytiva).

---

**Supplementary Table S1.**
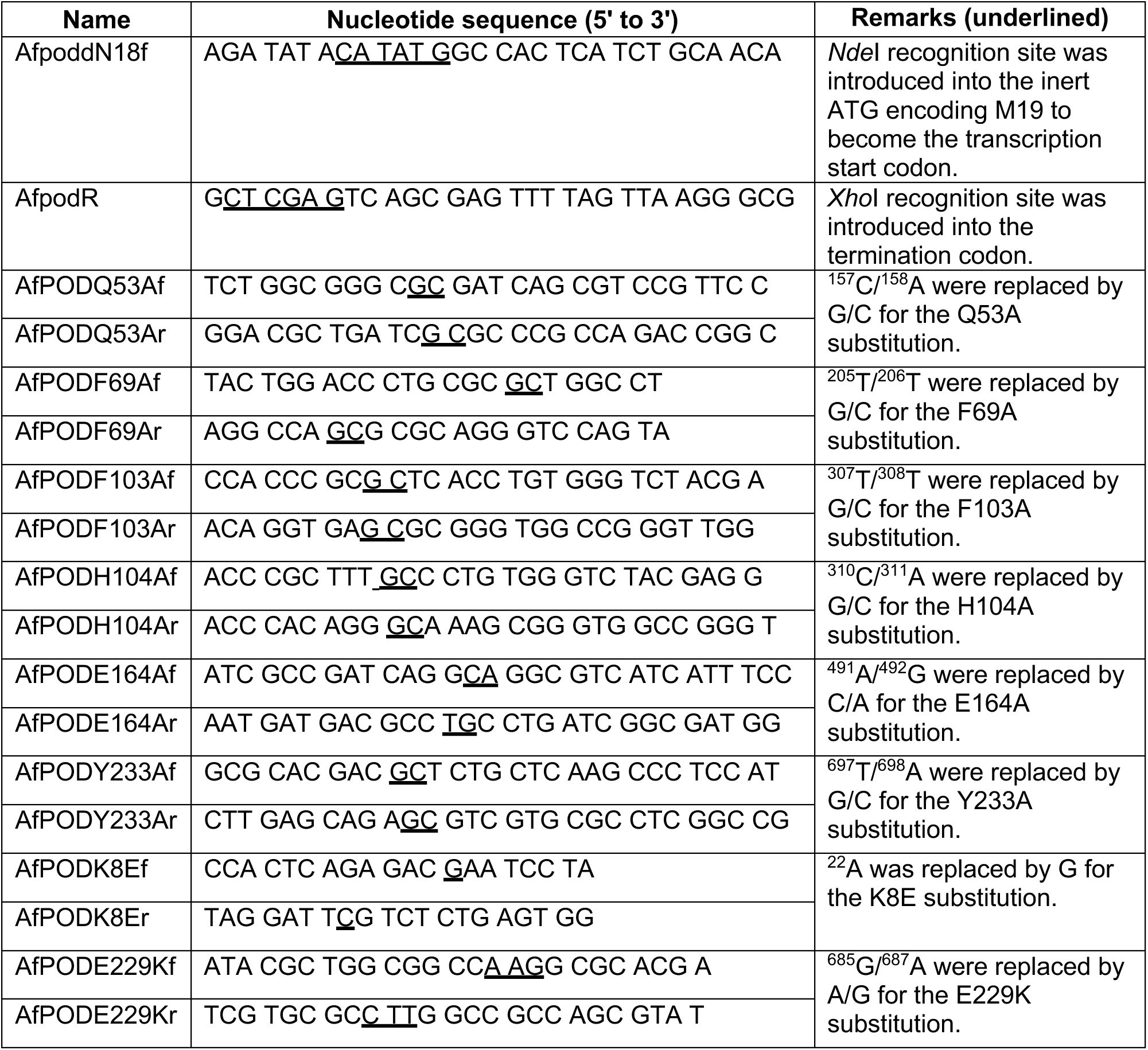
Primers used in this study.

**Supplementary Table S2.**
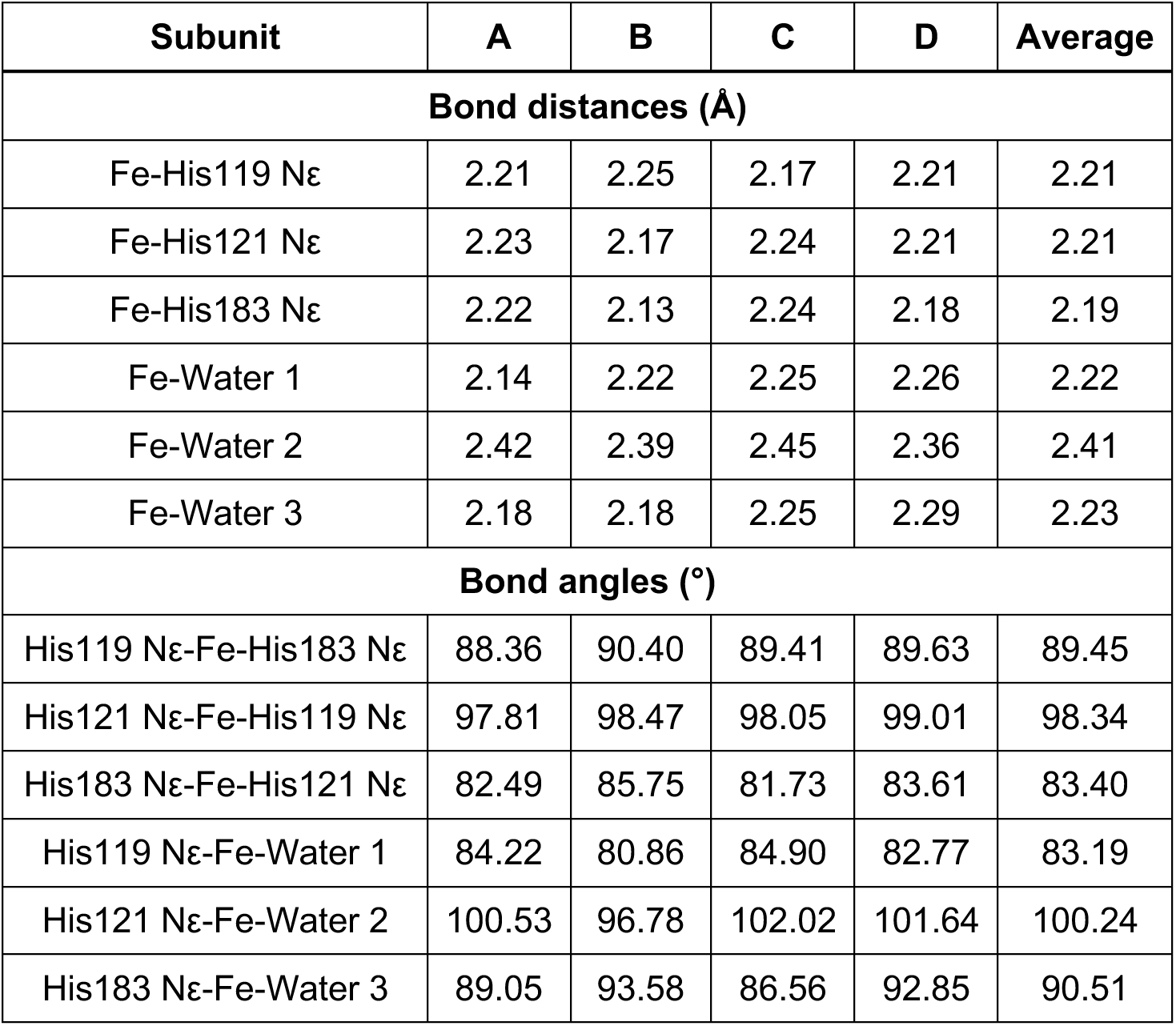
Geometry of the iron coordination sphere in the active site of AfPOD.

| Subunit | A | B | C | D | Average |
| --- | --- | --- | --- | --- | --- |
| Bond distances (Å) |  |  |  |  |  |
| Fe-His119 Nε | 2.21 | 2.25 | 2.17 | 2.21 | 2.21 |
| Fe-His121 Nε | 2.23 | 2.17 | 2.24 | 2.21 | 2.21 |
| Fe-His183 Nε | 2.22 | 2.13 | 2.24 | 2.18 | 2.19 |
| Fe-Water 1 | 2.14 | 2.22 | 2.25 | 2.26 | 2.22 |
| Fe-Water 2 | 2.42 | 2.39 | 2.45 | 2.36 | 2.41 |
| Fe-Water 3 | 2.18 | 2.18 | 2.25 | 2.29 | 2.23 |
| Bond angles (°) |  |  |  |  |  |
| His119 Nε-Fe-His183 Nε | 88.36 | 90.40 | 89.41 | 89.63 | 89.45 |
| His121 Nε-Fe-His119 Nε | 97.81 | 98.47 | 98.05 | 99.01 | 98.34 |
| His183 Nε-Fe-His121 Nε | 82.49 | 85.75 | 81.73 | 83.61 | 83.40 |
| His119 Nε-Fe-Water 1 | 84.22 | 80.86 | 84.90 | 82.77 | 83.19 |
| His121 Nε-Fe-Water 2 | 100.53 | 96.78 | 102.02 | 101.64 | 100.24 |
| His183 Nε-Fe-Water 3 | 89.05 | 93.58 | 86.56 | 92.85 | 90.51 |

**Supplementary Table S3.** List of docking results by Ginina.

| model | affinity<br>(kcal/mol) | intramol<br>(kcal/mol) | CNN<br>pose score | CNN<br>affinity |
| --- | --- | --- | --- | --- |
| 1 | -4.31 | -0.23 | 0.6880 | 4.192 |
| 2 | -3.90 | -0.23 | 0.6647 | 3.959 |
| 3 | -4.03 | -0.22 | 0.6381 | 3.932 |
| 4 | -4.05 | -0.24 | 0.6347 | 3.822 |
| 5 | -3.50 | -0.44 | 0.6210 | 3.752 |
| 6 | -4.04 | -0.22 | 0.5956 | 3.845 |
| 7 | -3.44 | -0.23 | 0.5825 | 3.550 |
| 8 | -3.34 | -0.20 | 0.5806 | 3.503 |
| 9 | -3.36 | -0.21 | 0.5793 | 3.554 |
| 10 | -3.82 | -0.24 | 0.5310 | 3.742 |
| 11 | -4.03 | -0.23 | 0.5310 | 3.321 |
| 12 | -3.54 | -0.07 | 0.5185 | 3.696 |
| 13 | -3.25 | -0.24 | 0.5154 | 3.376 |
| 14 | -4.16 | -0.22 | 0.5128 | 3.456 |
| 15 | -3.57 | -0.24 | 0.4871 | 3.709 |
| 16 | -3.59 | -0.23 | 0.4850 | 3.680 |
| 17 | -3.68 | -0.23 | 0.4599 | 3.250 |
| 18 | -4.32 | -0.21 | 0.4327 | 2.953 |
| 19 | -3.64 | -0.20 | 0.4172 | 3.404 |
| 20 | -4.58 | -0.23 | 0.4155 | 2.989 |

**Supplementary Table S4.**
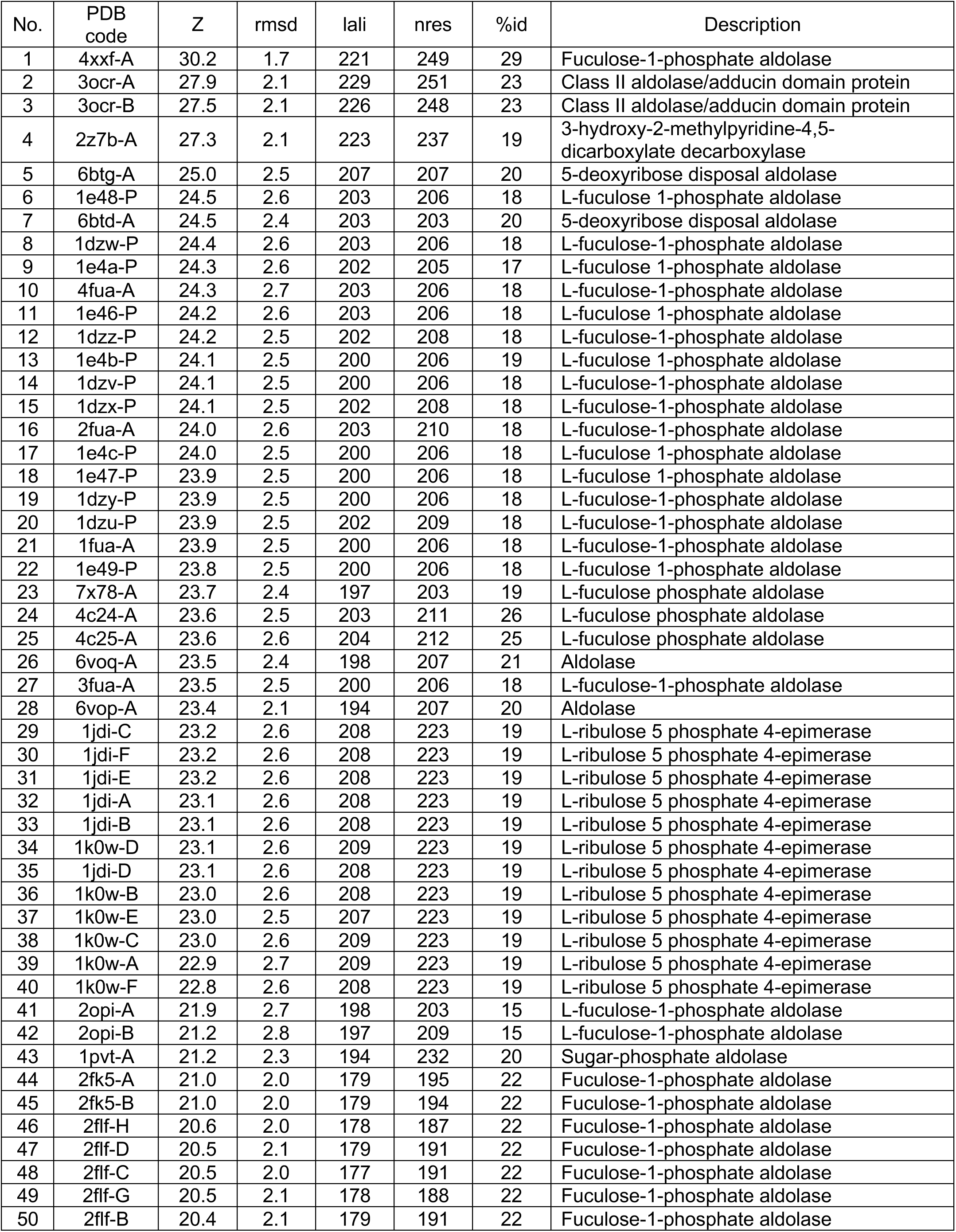
Structural similarity search by using the DALI server.

**Fig. S1.**
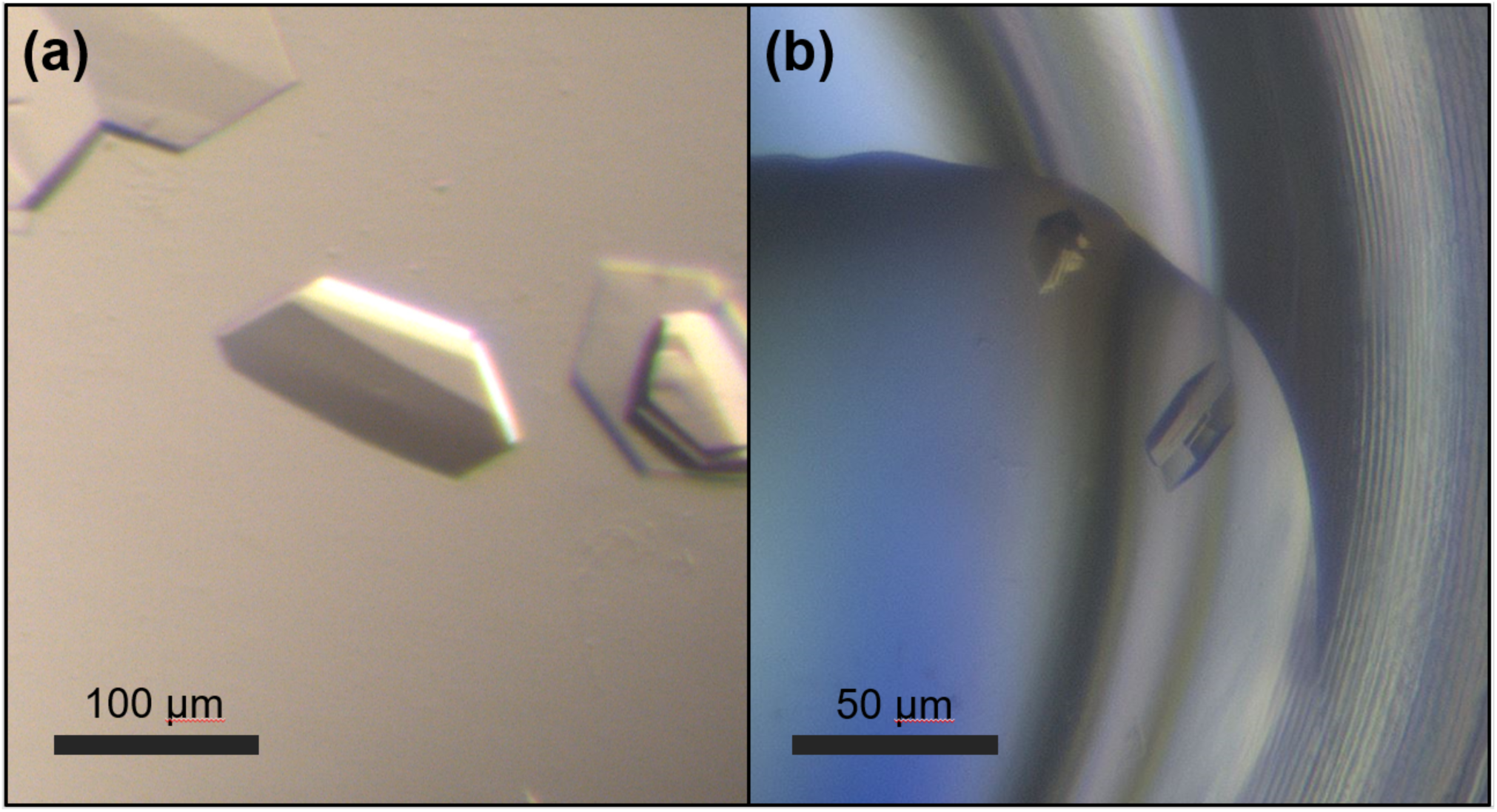
Crystals of (a) AfPOD and (b) AfPODdN18.

**Fig. S2.**
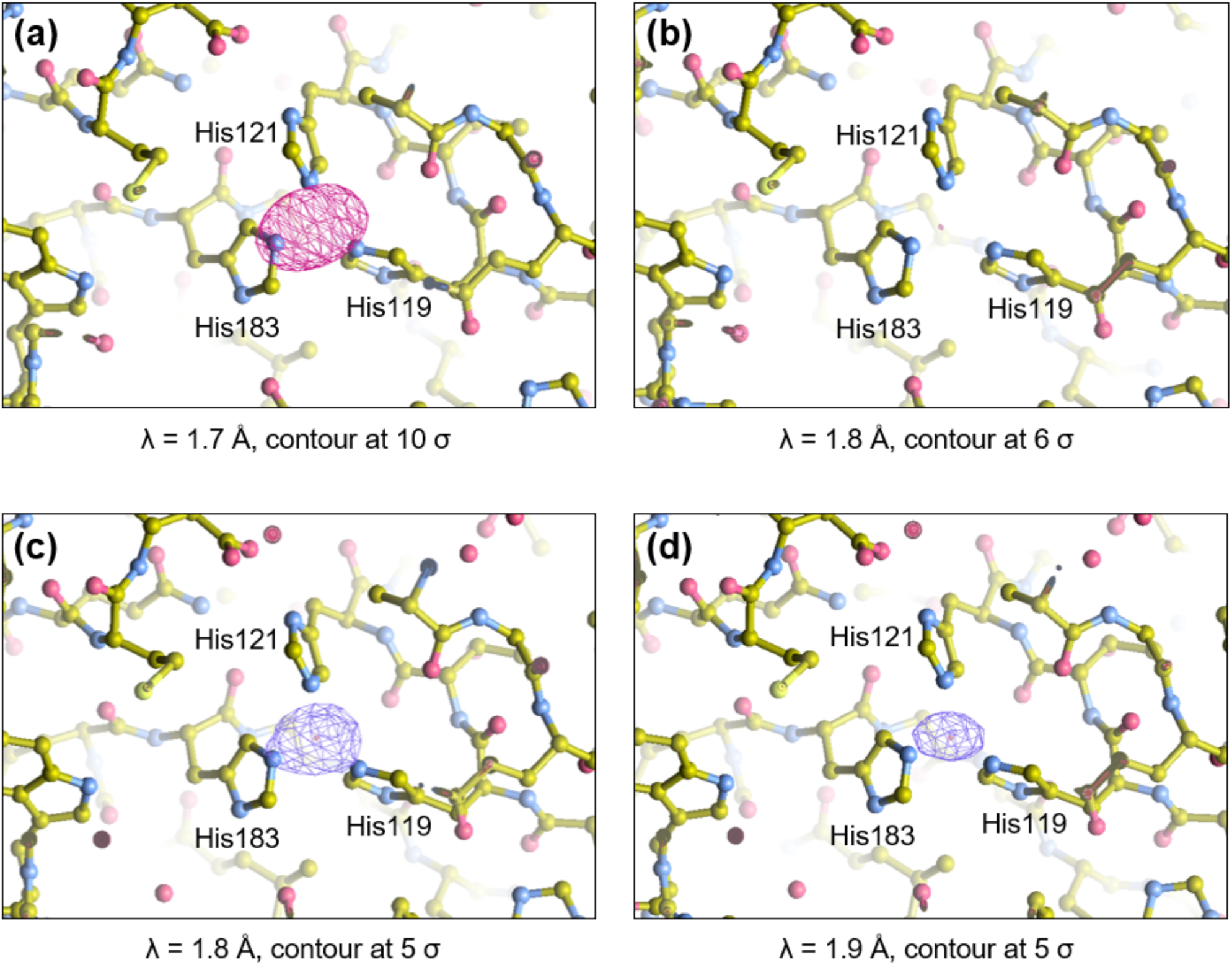
Identification of metal species at the active sites of AfPOD and AfPODdN18. Anomalous difference Fourier map of AfPOD obtained by using the diffraction data collected at (**a**) 1.7 Å and (**b**) 1.8 Å. Anomalous difference Fourier map of AfPODdN18 obtained by using the diffraction data collected at (**c**) 1.8 Å and (**d**) 1.9 Å.

**Fig. S3.**
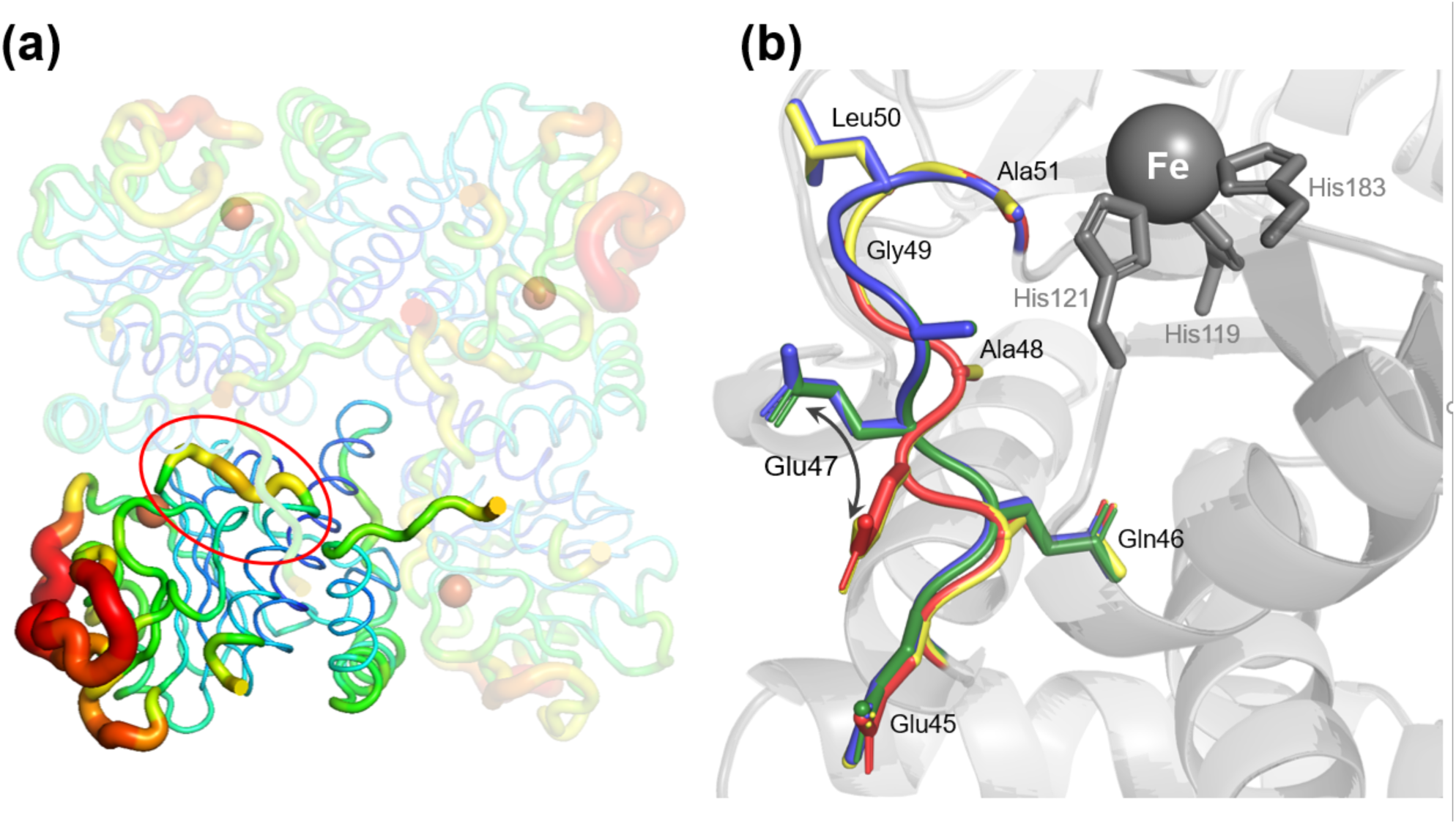
The mobile loop of AfPOD. (**a**) B-factor putty representation of the AfPOD structure. An α1-β1 loop (Glu45∼Ala51) near the iron center is circled in red. (**b**) The mobile loops of the AfPOD subunits A (blue), B (yellow), C (green), and D (red) have been superimposed.

**Fig. S4.**
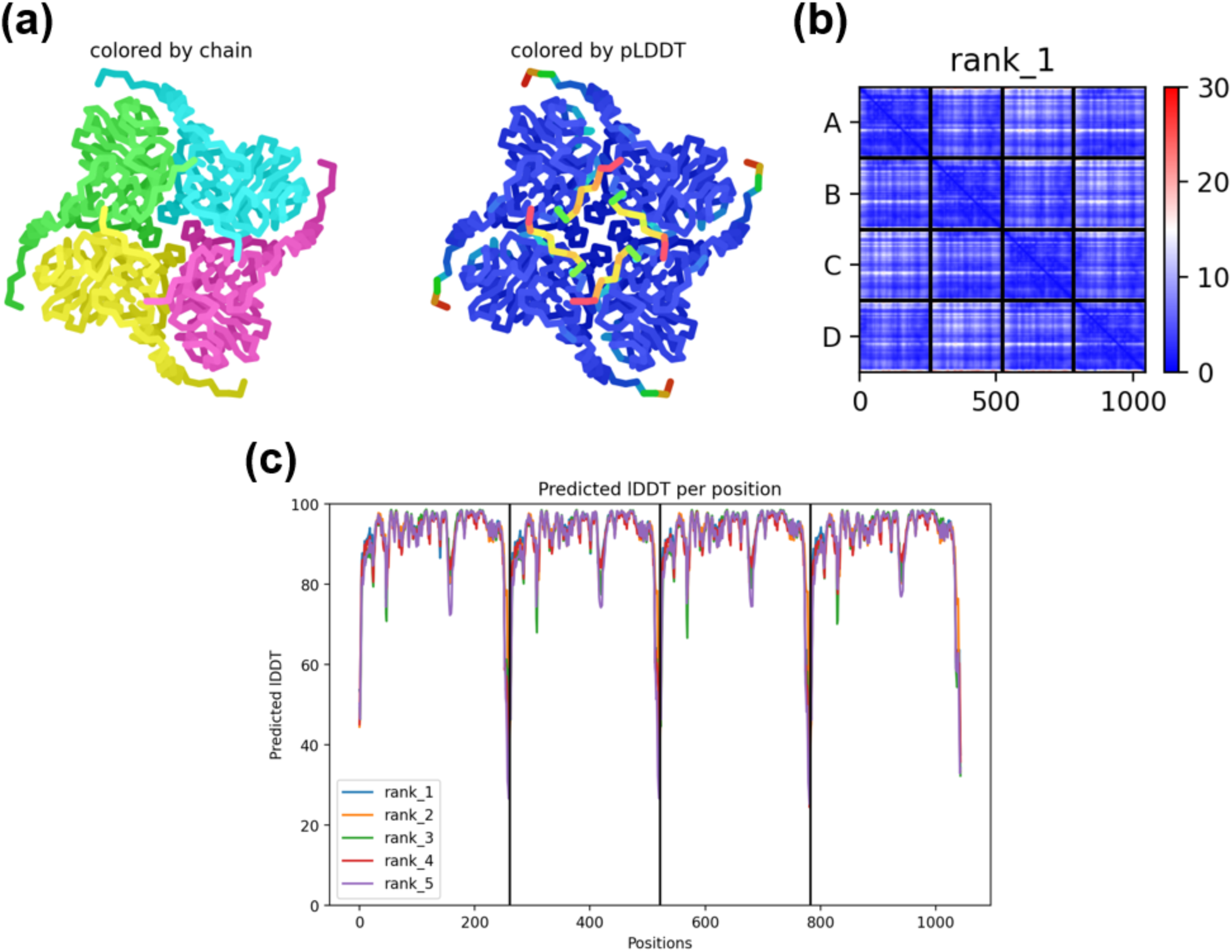
Confidence metrics for the predicted structure of AfPOD. (**a**) The predicted rank_1 structural model of AfPOD homotetramer colored by chain and by pLDDT (predicted Local Distance Difference Test). (**b**) Prediction aligned error (PAE) score for the rank_1 model. This score shows the calculated error of the predicted distance for each pair of residues. Both axes indicate the position of the each amino acid. The uncertainty in the predicted distance of two amino acids is color-coded from blue (0 Å) to red (30 Å), as shown in the right bar. (**c**) The pLDDT score per position for the five models generated by AlphaFold2 for the AfPOD homotetramer. The amino acid position is plotted against the pLDDT.

**Fig. S5.**
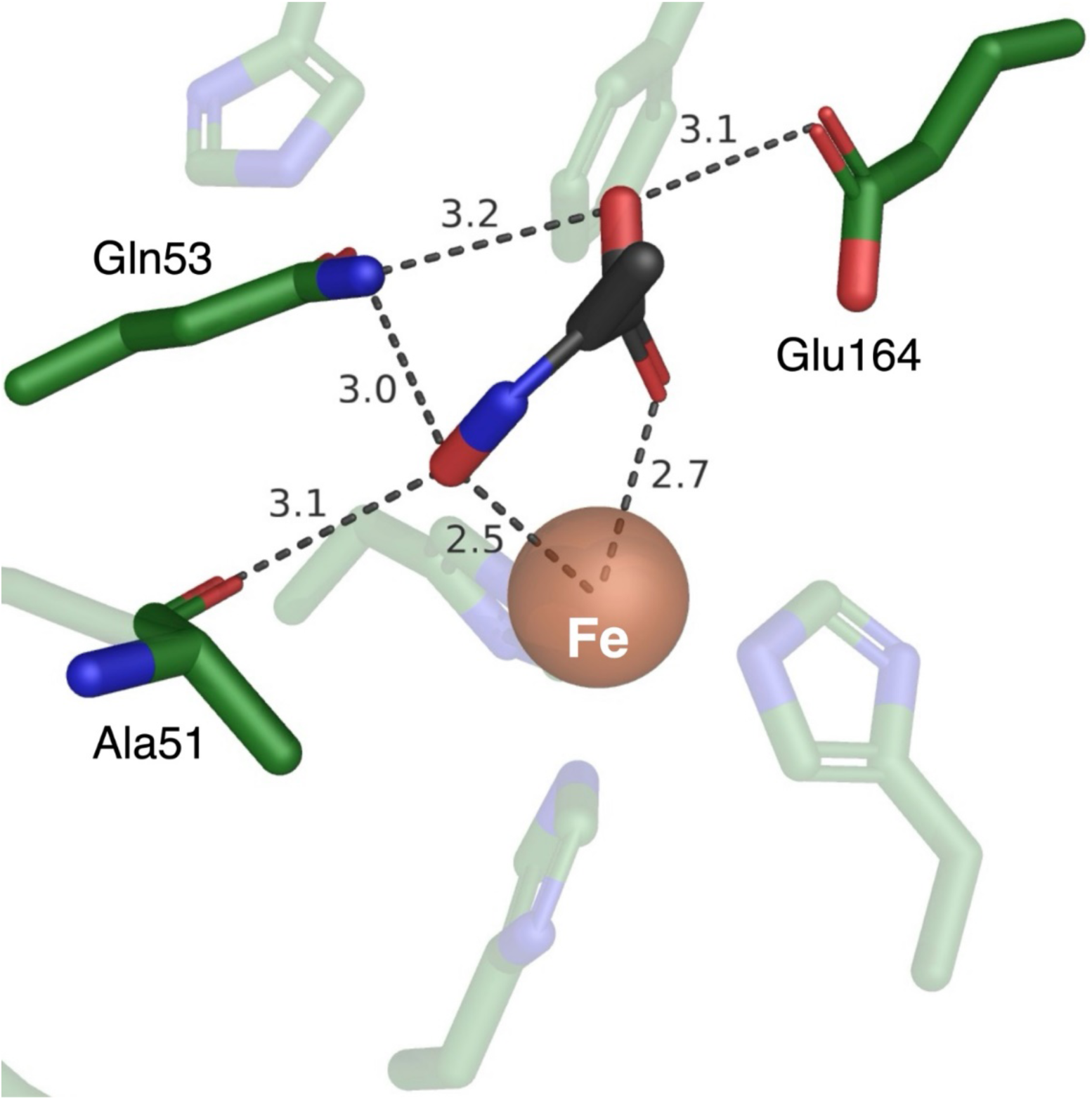
The docking structure with the highest pose score (model 1) of the active site of AfPOD with pyruvic oxime.

## REFERENCES

1. Wendeborn S. 2020. The chemistry, biology, and modulation of ammonium nitrification in soil. Angew Chem Int Ed Engl 59:2182–2202.

2. Rahimi S, Modin O, Mijakovic I. 2020. Technologies for biological removal and recovery of nitrogen from wastewater. Biotechnol Adv 43:107570.

3. Stein LY. 2011. Heterotrophic nitrification and nitrifier denitrification, p 95–114. *In* Ward BB, Arp DJ, Klotz MG (ed), Nitrification. ASM Press, Washington, DC.

4. Papen H, von Berg R. 1998. A Most Probable Number method (MPN) for the estimation of cell numbers of heterotrophic nitrifying bacteria in soil. Plant Soil 199:123–130.

5. Jensen HLY. 1951. Nitrification of oxime compounds by heterotrophic bacteria. J Gen Microbiol 5:360–368.

6. Castignetti D, Hollocher TC. 1982. Nitrogen redox metabolism of a heterotrophic, nitrifying-denitrifying *Alcaligenes* sp. from soil. Appl Environ Microbiol 44:923–928.

7. Castignetti D, Hollocher TC. 1984. Heterotrophic nitrification among denitrifiers. Appl Environ Microbiol 47:620–623.

8. Ono Y, Makino N, Hoshino Y, Shoji K, Yamanaka T. 1996. An iron dioxygenase from *Alcaligenes faecalis* catalyzing the oxidation of pyruvic oxime to nitrite. FEMS Microbiol Lett 139:103–108.

9. Tsujino S, Uematsu C, Dohra H, Fujiwara T. 2017. Pyruvic oxime dioxygenase from heterotrophic nitrifier *Alcaligenes faecalis* is a nonheme Fe(II)-dependent enzyme homologous to class II aldolase. Sci Rep 7:39991.

10. Tsujino S, Dohra H, Fujiwara T. 2021. Gene expression analysis of *Alcaligenes faecalis* during induction of heterotrophic nitrification. Sci Rep 11:23105.

11. Tsujino S, Masuda R, Shimizu Y, Azuma Y, Kanada Y, Fujiwara T. 2023. Phylogenetic diversity, distribution, and gene structure of the pyruvic oxime dioxygenase involved in heterotrophic nitrification. Antonie Van Leeuwenhoek 116:1037–1055.

12. Schägger H, von Jagow G. 1987. Tricine-sodium dodecyl sulfate-polyacrylamide gel electrophoresis for the separation of proteins in the range from 1 to 100 kDa. Anal Biochem 166:368–379.

13. Kato R, Hiraki M, Yamada Y, Tanabe M, Senda T. 2021. A fully automated crystallization apparatus for small protein quantities. Acta Crystallogr F 77:29–36.

14. Senda M, Senda T. 2018. Anaerobic crystallization of proteins. Biophys Rev 10:183–189.

15. Kabsch W. 2010. XDS. Acta Crystallogr D 66:125–132.

16. Skubák P, Pannu NS. 2013. Automatic protein structure solution from weak X-ray data. Nat Commun 4:2777.

17. Liebschner D, Yamada Y, Matsugaki N, Senda M, Senda T. 2016. On the influence of crystal size and wavelength on native SAD phasing. Acta Crystallogr D 7:728–741.

18. Vonrhein C, Flensburg C, Keller P, Sharff A, Smart O, Paciorek W, Womack T, Bricogne G. 2011. Data processing and analysis with the autoPROC toolbox. Acta Crystallogr D 6:293–302.

19. Vagin A, Teplyakov A. 1997. MOLREP: An automated program for molecular replacement. J Appl Cryst 30:1022–1025.

20. Emsley P, Cowtan K. 2004. Coot: model-building tools for molecular graphics. Acta Crystallogr D 60: 2126–2132.

21. Murshudov GN, Skubák P, Lebedev AA, Pannu NS, Steiner RA, Nicholls RA, Winn MD, Long F, Vagin AA. 2011. REFMAC5 for the refinement of macromolecular crystal structures. Acta Crystallogr. D 67:355–367.

22. Bricogne G, Blanc E, Brandl M, Flensburg C, Keller P, Paciorek W, Roversi P, Sharff A, Smart O, Vonrhein C, Womack T. 2022. BUSTER version 2.10.4. Cambridge, United Kingdom: Global Phasing Ltd.

23. Madeira F, Pearce M, Tivey ARN, Basutkar P, Lee J, Edbali O, Madhusoodanan N, Kolesnikov A, Lopez R. 2022. Search and sequence analysis tools services from EMBL-EBI in 2022. Nucleic Acids Res 50:W276–279.

24. Kumar S, Stecher G, Li M, Knyaz C, Tamura K. 2018. MEGA X: Molecular evolutionary genetics analysis across computing platforms. Mol Biol Evol 35:1547–1549.

25. Ashkenazy H, Abadi S, Martz E, Chay O, Mayrose I, Pupko T, Ben-Tal N. 2016. ConSurf 2016: an improved methodology to estimate and visualize evolutionary conservation in macromolecules. Nucleic Acids Res 44:W344–W350.

26. Holm L. 2022. Dali server: structural unification of protein families. Nucleic Acids Res 50:W210–W215.

27. Jumper J, Evans R, Pritzel A, Green T, Figurnov M, Ronneberger O, Tunyasuvunakool K, Bates R, Žídek A, Potapenko A, Bridgland A, Meyer C, Kohl SAA, Ballard AJ, Cowie A, Romera-Paredes B, Nikolov S, Jain R, Adler J, Back T, Petersen S, Reiman D, Clancy E, Zielinski M, Steinegger M, Pacholska M, Berghammer T, Bodenstein S, Silver D, Vinyals O, Senior AW, Kavukcuoglu K, Kohli P, Hassabis D. 2021. Highly accurate protein structure prediction with AlphaFold. Nature 596:583–589.

28. Mirdita M, Schütze K, Moriwaki Y, Heo L, Ovchinnikov S, Steinegger M. 2022. ColabFold: making protein folding accessible to all. Nat Methods 19:679–682.

29. Nicholas DJD, Nason A. 1957. Determination of nitrate and nitrite. Meth Enzymol 3:981–984.

30. Quastel JH, Scholefield PG, Stevenson JW. 1952. Oxidation of pyruvic acid oxime by soil organisms. Biochem J 51:278–286.

31. McNutt AT, Francoeur P, Aggarwal R, Masuda T, Meli R, Ragoza M, Sunseri J, Koes DR. 2021 GNINA 1.0: molecular docking with deep learning. J Cheminform 13:43.

32. Dreyer MK, Schulz GE. 1996. Catalytic mechanism of the metal-dependent fuculose aldolase from *Escherichia coli* as derived from the structure. J Mol Biol 259:458–466.

33. Krissinel E, Henrick K. 2007. Inference of macromolecular assemblies from crystalline state. J Mol Biol 372:774–797.

34. Wilson CJ, Choy WY, Karttunen M. 2022. AlphaFold2: A role for disordered protein/region prediction? Int J Mol Sci 23:4591.

35. Wang W, Liang AD, Lippard SJ. 2015. Coupling oxygen consumption with hydrocarbon oxidation in bacterial multicomponent monooxygenases. Acc Chem Res 48:2632–2639.

36. Irmisch S, Clavijo McCormick A, Günther J, Schmidt A, Boeckler GA, Gershenzon J, Unsicker SB, Köllner TG. 2014. Herbivore-induced poplar cytochrome P450 enzymes of the CYP71 family convert aldoximes to nitriles which repel a generalist caterpillar. Plant J 80:1095–1107.

37. Beaudoin GAW, Li Q, Folz J, Fiehn O, Goodsell JL, Angerhofer A, Bruner SD, Hanson AD. 2018. Salvage of the 5-deoxyribose byproduct of radical SAM enzymes. Nat Commun 9:3105.

38. Luo Y, Samuel J, Mosimann SC, Lee JE, Tanner ME, Strynadka NC. 2001. The structure of L-ribulose-5-phosphate 4-epimerase: an aldolase-like platform for epimerization. Biochemistry 40:14763–14771.

39. Kroemer M, Schulz GE. 2002. The structure of L-rhamnulose-1-phosphate aldolase (class II) solved by low-resolution SIR phasing and 20-fold NCS averaging. Acta Crystallogr D 58:824–832.

40. Mukherjee T, McCulloch KM, Ealick SE, Begley TP. 2007. Gene identification and structural characterization of the pyridoxal 5’-phosphate degradative protein 3-hydroxy-2-methylpyridine-4,5-dicarboxylate decarboxylase from *Mesorhizobium loti* MAFF303099. Biochemistry 46:13606–13615.

41. de Visser SP, Straganz GD. 2009. Why do cysteine dioxygenase enzymes contain a 3-His ligand motif rather than a 2His/1Asp motif like most nonheme dioxygenases? J Phys Chem A 113:1835–1846.

42. Senda T, Sugiyama K, Narita H, Yamamoto T, Kimbara K, Fukuda M, Sato M, Yano K, Mitsui Y. 1996. Three-dimensional structures of free form and two substrate complexes of an extradiol ring-cleavage type dioxygenase, the BphC enzyme from *Pseudomonas* sp. strain KKS102. J Mol Biol 255:735–752.

43. Dreyer MK, Schulz GE. 1996. Refined high-resolution structure of the metal-ion dependent l-fuculose-1-phosphate aldolase (class II) from *Escherichia coli*. Acta Crystallogr D 52:1082–1091.

44. Joerger AC, Gosse C, Fessner WD, Schulz GE. 2000. Catalytic action of fuculose 1-phosphate aldolase (class II) as derived from structure-directed mutagenesis. Biochemistry 39:6033–6041.

45. Joerger AC, Mueller-Dieckmann C, Schulz GE. 2000. Structures of l-fuculose-1-phosphate aldolase mutants outlining motions during catalysis. J Mol Biol 303:531–543.

## REFERENCES

1. Tsujino S, Uematsu C, Dohra H, Fujiwara T. 2017. Pyruvic oxime dioxygenase from heterotrophic nitrifier *Alcaligenes faecalis* is a nonheme Fe(II)-dependent enzyme homologous to class II aldolase. Sci Rep 7:39991.

2. Sambrook J, Russell DW. 2001. Molecular cloning: a laboratory manual, 3rd edn. Cold Spring Harbor Laboratory Press, New York.

